# GR chaperone cycle mechanism revealed by cryo-EM: inactivation of GR by GR:Hsp90:Hsp70:Hop client-loading complex

**DOI:** 10.1101/2020.11.05.370247

**Authors:** Ray Yu-Ruei Wang, Chari M. Noddings, Elaine Kirschke, Alexander G. Myasnikov, Jill L. Johnson, David A. Agard

## Abstract

Maintaining a healthy proteome is fundamental for organism survival^1,2^. Integral to this are Hsp90 and Hsp70 molecular chaperones that together facilitate the folding, remodeling and maturation of Hsp90’s many “client” proteins^3–7^. The glucocorticoid receptor (GR) is a model client strictly dependent upon Hsp90/Hsp70 for activity^8–13^. Chaperoning GR involves a cycle of inactivation by Hsp70, formation of an inactive GR:Hsp90:Hsp70:Hop “loading” complex, conversion to an active GR:Hsp90:p23 “maturation” complex, and subsequent GR release^14^. Unfortunately, a molecular understanding of this intricate chaperone cycle is lacking for any client. Here, we report the cryo-EM structure of the GR loading complex, in which Hsp70 loads GR onto Hsp90, revealing the molecular basis of direct Hsp90/Hsp70 coordination. The structure reveals two Hsp70s—one delivering GR and the other scaffolding Hop. Unexpectedly, the Hop cochaperone interacts with all components of the complex including GR, poising Hsp90 for subsequent ATP hydrolysis. GR is partially unfolded and recognized via an extended binding pocket composed of Hsp90, Hsp70 and Hop, revealing the mechanism of GR loading and inactivation. Together with the GR maturation complex (Noddings et al., 2020), we present the first complete molecular mechanism of chaperone-dependent client remodeling, establishing general principles of client recognition, inhibition, transfer and activation.

## Main

The highly abundant and conserved Hsp90 and Hsp70 molecular chaperones are essential for proteome maintenance. Hsp70 recognizes virtually all unfolded/misfolded proteins, and generally functions early in protein folding^15^. By contrast, Hsp90 typically functions later^11^, targeting a select set of “clients”^7,16^. Despite the differences, Hsp90 and Hsp70 share clients that are highly enriched for signaling and regulatory proteins^4,16,17^, making both chaperones important pharmaceutical targets for cancer^18,19^ and neurodegenerative diseases^20–22^. Both chaperones are dynamic molecular machines with complex ATP-dependent conformational cycles that drive client binding/release. Hsp70 uses its N-terminal nucleotide-binding domain (Hsp70_NBD_) to allosterically regulate its C-terminal substrate-binding domain (Hsp70_SBD_), comprising a β-sandwich core (Hsp70_SBD-β_) and an α-helical lid (Hsp70_SBD-α_)^15,23–26^. In the weak client binding ATP-bound “open” state (Hsp70^ATP^), both the Hsp70_SBD-α_ and Hsp70_SBD-β_ dock onto the Hsp70_NBD_^27^. In the ADP state (Hsp70^ADP^), the Hsp70_NBD_ and Hsp70_SBD_ subdomains separate, resulting in a high-affinity client-binding state^28^. Hsp90 constitutively dimerizes through its C-terminal domain (Hsp90_CTD_) and cycles through open and closed conformations acting as a molecular clamp^5,29^. In the nucleotide-free state (Hsp90^Apo^), Hsp90 populates a variety of open conformations^29–31^, whereas ATP binding (Hsp90^ATP^) drives clamp closure via secondary dimerization of the N-terminal domains (Hsp90_NTD_)^32^. Clamp closure activates Hsp90 for ATP hydrolysis and is a rate-limiting^33^ process requiring Hsp90_NTD_ rotation, N-terminal helix rotation and ATP-binding pocket lid closure. Unlike Hsp70, Hsp90 can engage clients independent of nucleotide state via the middle domain^34,35^ (Hsp90_MD_) and the amphipathic helix-hairpins^36^ (Hsp90_amphi-α) on_ the Hsp90_CTD_.

The glucocorticoid receptor (GR) is a steroid hormone-activated transcription factor that constitutively depends on Hsp90 to function^10^. Building on the pioneering work of Pratt and Toft^8,12,13^, we previously reconstituted GR’s Hsp90 dependence using an *in vitro* system^14^, establishing a 4-step cycle (Fig. 1a) starting with active GR ligand-binding domain (hereafter GR for simplicity). Next, Hsp70 inactivates GR ligand binding, then co-chaperone Hop (Hsp90/Hsp70 organization protein) helps load Hsp70:GR onto Hsp90 forming the inactive “loading” complex (GR:Hsp90:Hsp70:Hop). Upon Hsp90 closure and ATP hydrolysis, Hsp70 and Hop are released, followed by the incorporation of p23, forming the GR:Hsp90:p23 “maturation” complex. In the maturation complex, GR is reactivated, indicating GR is conformationally remodeled during the transition. A similar pattern of Hsp70/Hsp90 functional antagonism has subsequently been shown for other clients^37–40^, supporting a general mechanism.

**Fig. 1.**
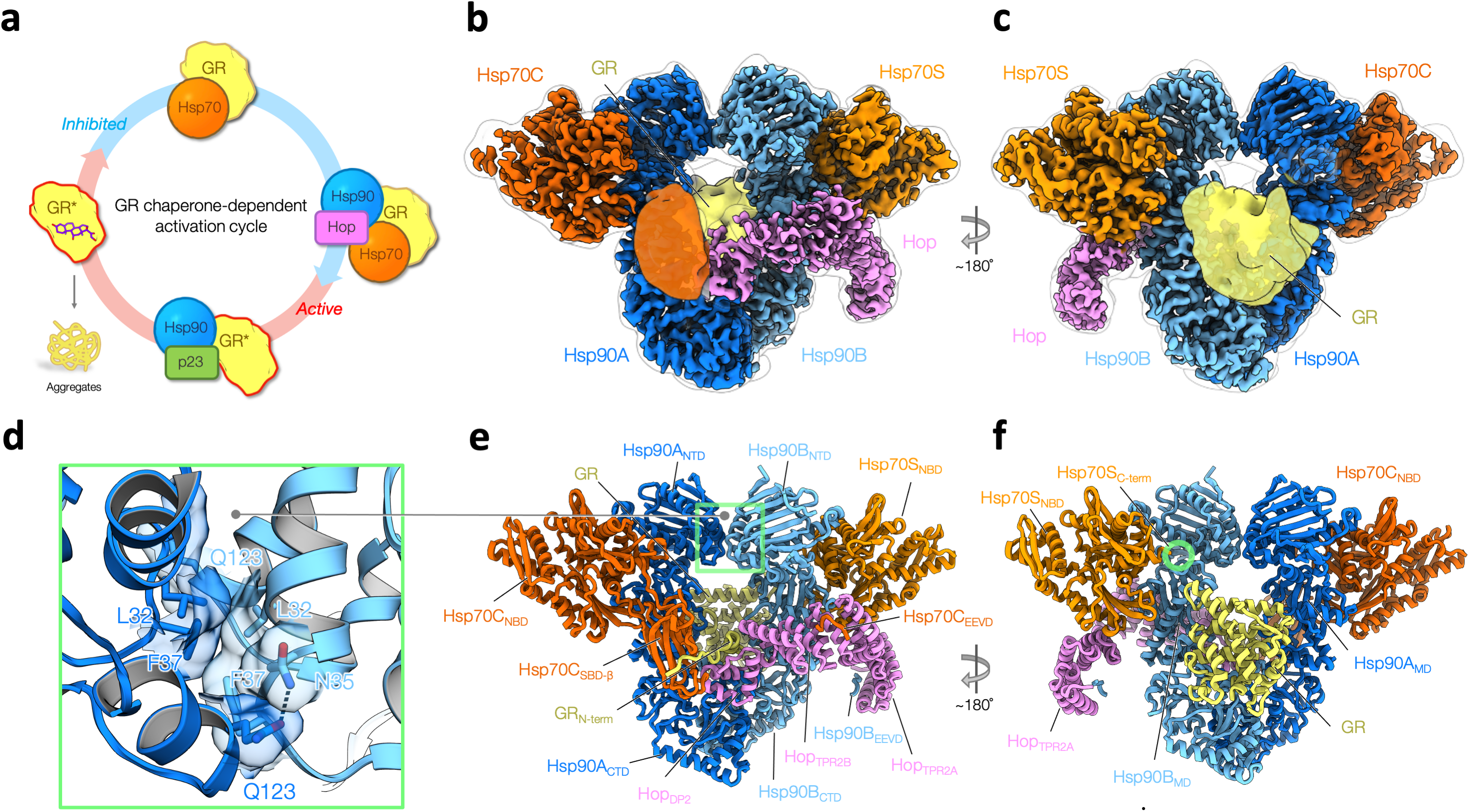
Overview of the GR-loading complex. **a,** GR activity is regulated by molecular chaperones in a constant cycle. Starting from active GR (left), which is aggregation-prone (lower left) in physiological conditions, Hsp70 protects and inhibits GR (top). Recruited by Hop, Hsp70 loads GR to Hsp90, forming the loading complex, in which GR remains inactive. Upon ATP hydrolysis on Hsp90, GR is reactivated in the maturation complex of GR:Hsp90:p23 (bottom) and is thereafter released to continue the cycle. **b,c**, Front (**b**) and back (**c**) views of a composite cryo-EM map of the GR:Hsp90:Hsp70:Hop complex. Densities of Hsp70C_SBD_ and the globular C-terminal GR (yellow) are taken from the low-pass-filtered map of the high-resolution reconstruction of the full complex. The subunit color code is used throughout. **d,** Close-up view of the novel dimerization interface of the symmetric Hsp90 dimer. The interface is composed of two molecular switches of Hsp90, the first helix and the lid motif. Dashed line depicts a polar interaction. **e,f,** Corresponding views of the atomic model of the GR-loading complex. Green circle indicates the C-terminus of Hsp70S.

Unfortunately, the molecular basis for almost all of this complex chaperone interplay remains unknown, with high-resolution structural studies hampered by the instability of clients and the highly dynamic nature of client:chaperone associations. Here, we report a high-resolution cryo-EM structure of the client-loading complex, providing much needed molecular insights into how Hsp90/Hsp70 coordinate their ATP cycles, how they are organized by Hop, and the molecular mechanisms underlying GR’s functional regulation by Hsp90/Hsp70.

## Results

### Structure determination and architecture of the client-loading complex

The client-loading complex was prepared by reconstitution using excess ADP to enhance client binding by Hsp70 and an ATP-binding deficient Hsp90 (Hsp90^D93N^)^41^ to stall the cycle at this intermediate step, followed by glutaraldehyde stabilization (Extended Data Fig. 1). A ~3.6Å resolution cryo-EM reconstruction was obtained from ~4 million particles using RELION^42^ (Extended Data Fig. 2–4, Materials and Methods). The resulting structure reveals an architecture drastically different than expected, with the Hsp90 dimer (Hsp90A/B) surrounded by Hop, GR and unexpectedly two Hsp70s (Hsp70“C” for client-loading and Hsp70“S” for scaffolding) (Fig. 1b,c,e,f). Hsp90 adopts a previously unseen “semi-closed” conformation, in which the Hsp90_NTD_s have rotated into an Hsp90^ATP^-like orientation but have not yet reached the fully-closed ATP state (Extended Data Fig. 5). The observed Hsp90_NTD_ orientation is stabilized by two Hsp70_NBD_s that bind symmetrically to the Hsp90_NTD_/Hsp90_MD_ interface of each Hsp90 protomer. Hop intimately interacts with each Hsp90 protomer, the two Hsp70s, and remarkably, a portion of GR. Although the two Hsp70_SBD_s are not visible in the high-resolution map, the Hsp70C_SBD-β_ subdomain becomes visible in low-pass filtered maps (Fig. 1b). Also seen in the filtered maps, GR is positioned on one side of the Hsp90 dimer (Fig. 1c and Extended Data Fig. 23b,c). Additionally, a lower-resolution map (~7Å) was reconstructed showing a loading complex that has lost Hsp70C but retains Hsp70S, Hop and GR (Extended Data Fig. 6). The observation of the two-Hsp70 and one-Hsp70 loading complexes populated in our sample is consistent with a previous study^43^.

### The nucleotide-regulated interplay between Hsp90 and Hsp70

Two major interfaces are formed in both of the nearly identical Hsp70_NBD_/Hsp90 protomer interactions (RMSD of 0.96Å, Extended Data Fig. 7a-c). In Interface I, the outer edge of the Hsp90_MD_ β-sheet inserts into the cleft formed by the Hsp70_NBD-IA_ and Hsp70_NBD-IIA_ subdomains (Fig. 2a, Extended Data Fig. 8a). Notably, in Hsp70^ATP^ this cleft binds the Hsp70 interdomain linker^24,27^ and also contributes to binding Hsp40’s J-domain^44^ (Extended Data Fig. 8b). Hence, the cleft is only available in Hsp70^ADP^. Interface I is tightly packed (479Å^2^ of buried surface area, BSA), and is stabilized by numerous polar interactions (Fig. 2c,d). This explains the reduced Hsp70 interaction and the growth defects and significant loss of GR and v-Src function caused by mutations of the central residue in this interface Hsp90^G333 (G313 in yHsp82/G309 in yHsc82)^ (Fig. 2d) and the 2.5–5 fold reduction in Hsp90/Hsp70 affinity observed for nearby salt bridge mutations^45–47^ (Fig. 2c).

**Fig. 2.**
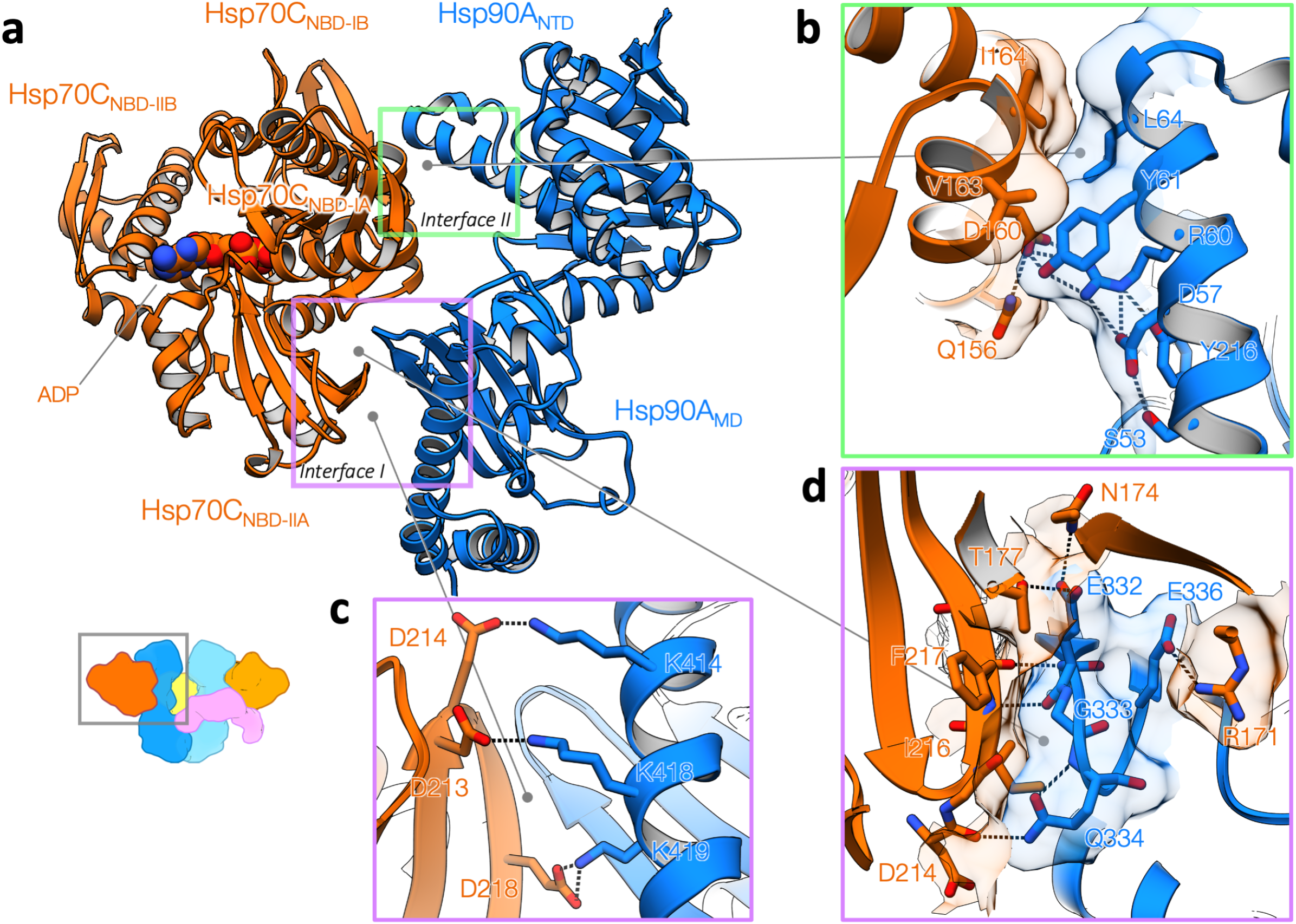
Molecular basis of Hsp90:Hsp70 interactions. **a**, Hsp90:Hsp70 interactions without the aid of Hop (Hsp90A:Hsp70C). In Interface I (purple rectangle), Hsp90 uses the outer edge of the Hsp90Ą_MD_ β-sheet to insert into a cleft formed by Hsp70_NBD-IA_ and Hsp70_NBD-IIA_ subdomains. The cleft is where the Hsp70 interdomain linker binds and serves as the allosteric center for Hsp70 NBD to regulate client binding in the Hsp70 SBD. In Interface II (green rectangle), the two ATPase domains Hsp90 and Hsp70 directly interact with each other. **b,** Close-up view of Interface II. Transparent surface and stick representations are shown for residues involved in the interactions. Dashed lines depict the network of polar interactions involved. **c,** Interface I is featured with conserved salt bridges (dashed lines). **d,** Detailed view of Interface II. A β-strand pairing (dashed lines) between the backbone atoms of Hsp90^E332^ and Hsp70^F217^, in which Hsp90^G333^ is closely packed with Hsp70^F217^.

Interface II (Fig. 2a,b, 280Å^2^ BSA) is stabilized by both hydrophobic (Hsp90^Y61,L64^:Hsp70^V163,I164^) and polar interactions (Hsp90^R60,Y61^: Hsp70^D160^). Importantly, it defines the Hsp90^ATP^-like position/orientation of the Hsp90_NTD_ with respect to the Hsp90_MD_, explaining the observation that Hsp70 accelerated Hsp90 ATPase activity^48^. Consistent with the significance of Interface II, mutation of the three Hsp90 interface residues (Hsp90^R60,Y61,L64^) showed marked yeast growth defects at 37°C^49^. Similar to Interface I mutations (yHsc82^G309S,E394K^), an Hsp90^R60^ mutation (yHsc82^R46G^) displayed reduced Hsp70 interaction, inviability at 37°C, and reduced v-Src activity (Extended Data Fig. 9). Lastly, sequence alignments of Hsp90/Hsp70 homologs/paralogues showed that Interface I & II residues are generally conserved, suggesting a universal Hsp70-Hsp90 binding strategy across species^47,50^ and organelles^51,52^ (Extended Data Fig. 10,11).

As expected from Hsp90^D93N^, both Hsp90_NTD_ ATP-binding pockets are empty with their lids open. The Hsp90A_NTD_ and Hsp90B_NTD_s closely resemble the structure of an apo Hsp90_NTD_ fragment^53^ (RMSD of 0.43 and 0.35Å to 3T0H, respectively, Extended Data Fig. 12a). The ATP pocket lid and the first α-helix form a novel dimerization interface (426Å^2^ BSA, Fig. 1d).The two Hsp70_NBD_s clearly have ADP bound and are similar to the ADP-bound Hsp70_NBD_ crystal structure^54^ (Cα-RMSD of 0.50Å (Hsp70C) and 0.53Å (Hsp70S) to 3AY9, respectively) (Fig. 2a, Extended Data Fig. 13a,b).

Coordination between the Hsp90/Hsp70 ATPase cycles is required for forming the loading complex. The Hsp90^ATP^ conformation is incompatible as closure of the Hsp90^ATP^ ATP pocket lid would clash with the Hsp70_NBD_ (Extended Data Fig. 12b,c). Thus, ATP binding to Hsp90 would be expected to accelerate loss of the bound Hsp70s. Furthermore, the Hsp70^ATP^ conformation is incompatible with the loading complex, as the entire Hsp70_SBD_ would clash with Hsp90_NTD_/Hsp90_MD_ (Extended Data Fig. 14a,b). For Hsp70 to reenter its ATP cycle, it must first leave Hsp90, thus nucleotide exchange on Hsp70 likely times its dissociation. Notably, in the complex Hsp70_NBD-IIA_ deviates from the crystal structure (Extended Data Fig. 13a,b), likely explaining the weak Hsp70 nucleotide exchange activity provided by Hsp90 during the GR-chaperoning cycle^55^. The canonical nucleotide exchange factor (NEF) binding sites^56^ on Hsp70_NBD-IIB_ remain available (Extended Data Fig. 15a,b), explaining how the NEF Bag-1 can accelerate GR maturation^55^.

### Hop interacts extensively with all components in the loading complex

The cochaperone Hop is well conserved in eukaryotes and facilitates GR maturation *in vivo*^57^ and *in vitro*^14^. Hop is thought to bring Hsp90 and Hsp70 together using its three TPR domains that bind the EEVD C-termini on both Hsp90 and Hsp70^58–60^. Despite using the full-length Hop construct, only three C-terminal domains (Hop_TPR2A_, Hop_TPR2B_, and Hop_DP2_) are observed (Fig. 1e,f). Importantly, these three domains are necessary and sufficient for full GR activation^61,62^. Hop wraps around much of the loading complex, with extensive interactions made by Hop_TPR2A_ and Hop_DP2_, demonstrating a far more integral role than anticipated (Fig. 3a,c).

**Fig. 3.**
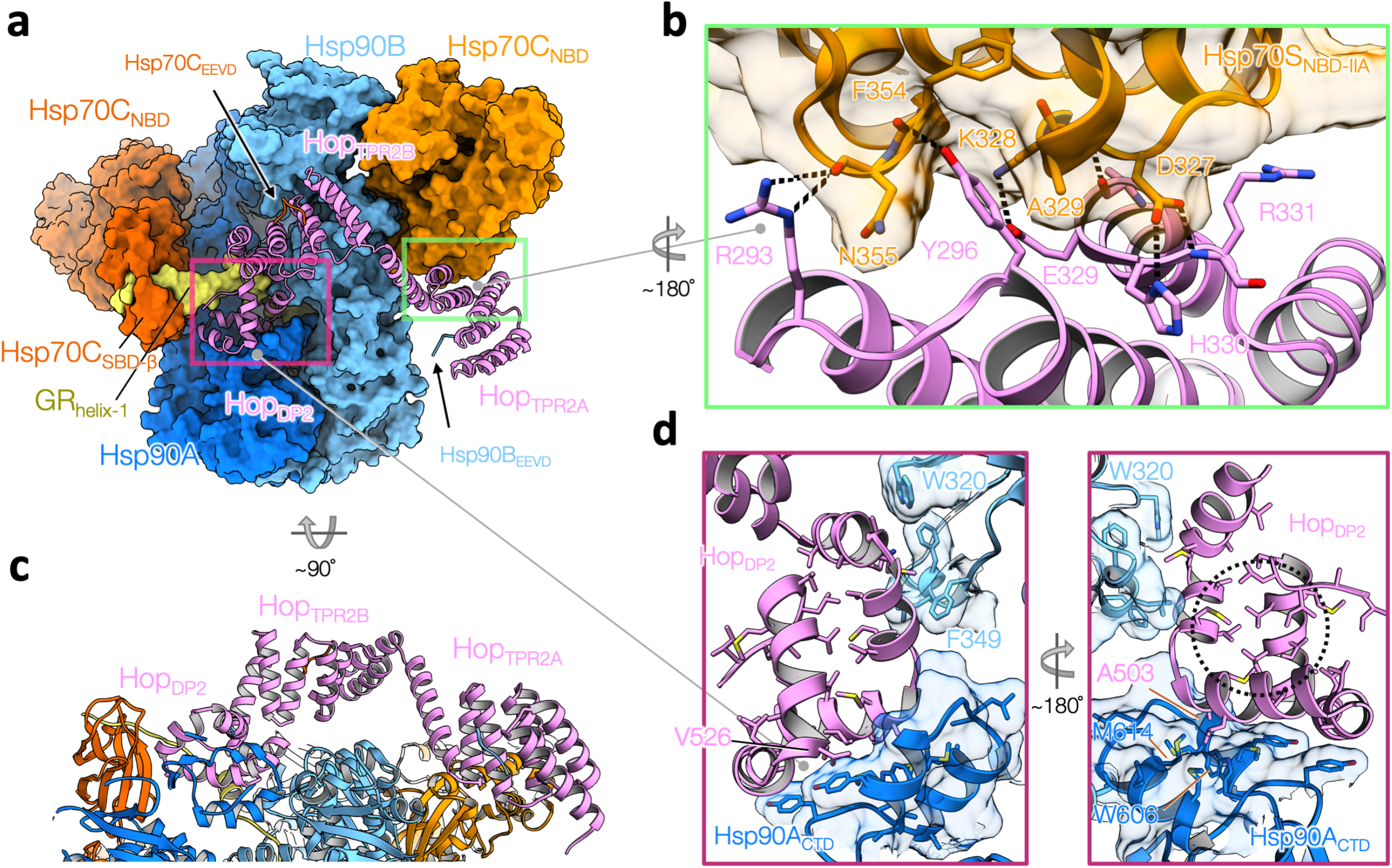
Hop interacts intimately with all components in the loading complex. **a,** Hop, shown in ribbon representation, uses its three C-terminal domains (Hop_TPR2A_, Hop_TRR2B_ and Hop_DP2_) to interact with Hsp90/Hsp70 beyond the EEVD binding. All the other components are shown in surface representation. The red rectangle highlights that Hop_DP2_ interacts with Hsp90A (dark blue), Hsp90B (light blue), Hsp70C_SBD_ (dark orange) and a portion of GR (yellow). The green rectangle highlights a novel interface formed by Hop_TPR2A_ and Hsp70C_NBD-IIA_. **b,** Close-up view of the novel Hop:Hsp70 interface with a 180-degree rotation from (a). Hop^Y296^ inserts into a cavity on Hsp70 (transparent surface representation), forming a hydrogen bond with the Hsp70^F354^ backbone atom. The interface also features many polar interactions depicted with dashed lines. **c,** A 90-degree rotation from the bottom view in (a) with ribbon model. No major interaction is observed between Hop_TPR2B_ and the loading complex. **d,** Left, Hop_DP2_ uses surface-exposed hydrophobic residues, shown in sticks, to interact with Hsp90’s client-binding motifs (shown in transparent surface and with hydrophobic residues in sticks)— the amphipathic helical hairpin from Hsp90A_CTD_ (dark blue) and Hsp90B^W320,F349^ (light blue). Right, a ~180-degree view from the left, Hop_DP2_ adopts a hand-like α-helical structure. The core of Hop_DP2_ is loosely packed with many hydrophobic residues exposed in the “palm” (black circle) of the “hand”. Note the GR-binding to Hop_DP2_ is not shown here.

The structure of Hop^TPR2A^-Hop^TRP2B^ closely matches the yeast crystal structure^61^ (Cα- RMSD of 1.47Å to 3UQ3, Extended Data Fig. 16a,c,d), including the conserved electrostatic network (Hop^Y354,R389,E385,K388^) that defines the unique inter-domain angle (Extended Data Fig. 17c-e). Focused maps revealed that the Hop_TPR2A_ and Hop_TPR2B_ are bound to the EEVD termini of Hsp90 and Hsp70, respectively (Extended Data Fig. 18c-e,19a-d). Although the density for the remaining Hsp70 and Hsp90 tails are missing, our structural modeling suggested the connectivity (Extended Data Fig. 20a,b). Unexpectedly, Hop_TPR2A_ and Hsp70S_NTD-IIA_ form a novel and extensive interface (578Å^2^ BSA) composed largely of polar interactions (Fig. 3a,b and Extended Data Fig. 18a-c,f). Notably, Hsp70S_NTD-IIA_ interacts with Hop and Hsp90 simultaneously, thereby rigidly positioning Hop with respect to Hsp90. Although Hop_TPR2B_ was in close proximity (~6 Å) to Hsp90_MD_ no major contacts were observed (Fig. 3c). However, the low-resolution Hsp90:Hop cryo-EM structure^63^ (Extended Data Fig. 21a) and previous studies^61,64^ show that Hop_TPR2B_ can make direct contacts with Hsp90_MD_. This suggests that Hop may first prepare Hsp90 for Hsp70 and client interaction, and subsequently rearrange upon Hsp70S_NBD_ binding (Extended Data Fig. 21b,c).

Hop_DP2_ makes extensive interactions with both Hsp90 protomers at Hsp90_ACTD_ and Hsp90B_MD_, thereby defining and maintaining the semi-closed Hsp90 conformation within the loading complex (Fig. 3a). Interestingly, conserved client-binding residues on Hsp90 are repurposed for Hop_DP2_ binding (Fig. 3d). Supporting our observations, Hsp90 mutations which would destabilize the Hop_DP2_-Hsp90A interface (yHsp82^W585T,M593T^ corresponding to hHsp90^W606,M614^) cause yeast growth defects^65^. Our Hop_DP2_ structure agrees well with the yeast NMR structure^61^ (Cα-RMSD of 1.13Å to 2LLW; Extended Data Fig. 16b), adopting a hand-like α-helical structure, with many of its core hydrophobic sidechains exposed in the “palm” of the “hand” (Fig. 3d). Importantly, this hydrophobic palm is continuous with the client binding surface provided by the lumen between the Hsp90 protomers, augmenting the Hsp90_Aamphi-α_ with a stronger and extensive hydrophobic binding capability.

### GR is unfolded, threaded through the Hsp90 lumen and bound by Hsp90, Hop and Hsp70

In the high-resolution map, a strand of density can be seen passing through the Hsp90 lumen (Extended Data Fig. 22b,c). In the low-pass filtered map, this density connects to the globular part of GR on one side of Hsp90 (Extended Data Fig. 23a,b). On the other side, a GR helix is surprisingly cradled in the Hop_DP2_ hydrophobic palm and the rest of the GR becomes a strand embedded in the Hsp70C_SBD-β_ substrate binding pocket(Fig. 4a-d and Extended Fig. 24d,25e). Thus, GR is partially unfolded and threaded through the Hsp90 lumen, reminiscent of how the CDK4 kinase was unfolded^66^ by the fully closed Hsp90^ATP^. To test this unexpected client:cochaperone interaction, we substituted Hop^Q512^ in Hop_DP2_ which is close to, but not directly interacting with, the GR helix with the photo-reactive unnatural amino acid p-benzoyl-phenylalanine (Extended Data Fig. 26b). In support of our structure, GR and Hop become photo-crosslinked (Extended Data Fig. 26a,c). Additionally, mutation of Hop^L508A (L553 in yeast, Sti1)^ which is in the hydrophobic palm that directly interacts with GR (Extended Data Fig. 27a), completely abrogated GR function *in vivo^61^.*

**Fig. 4.**
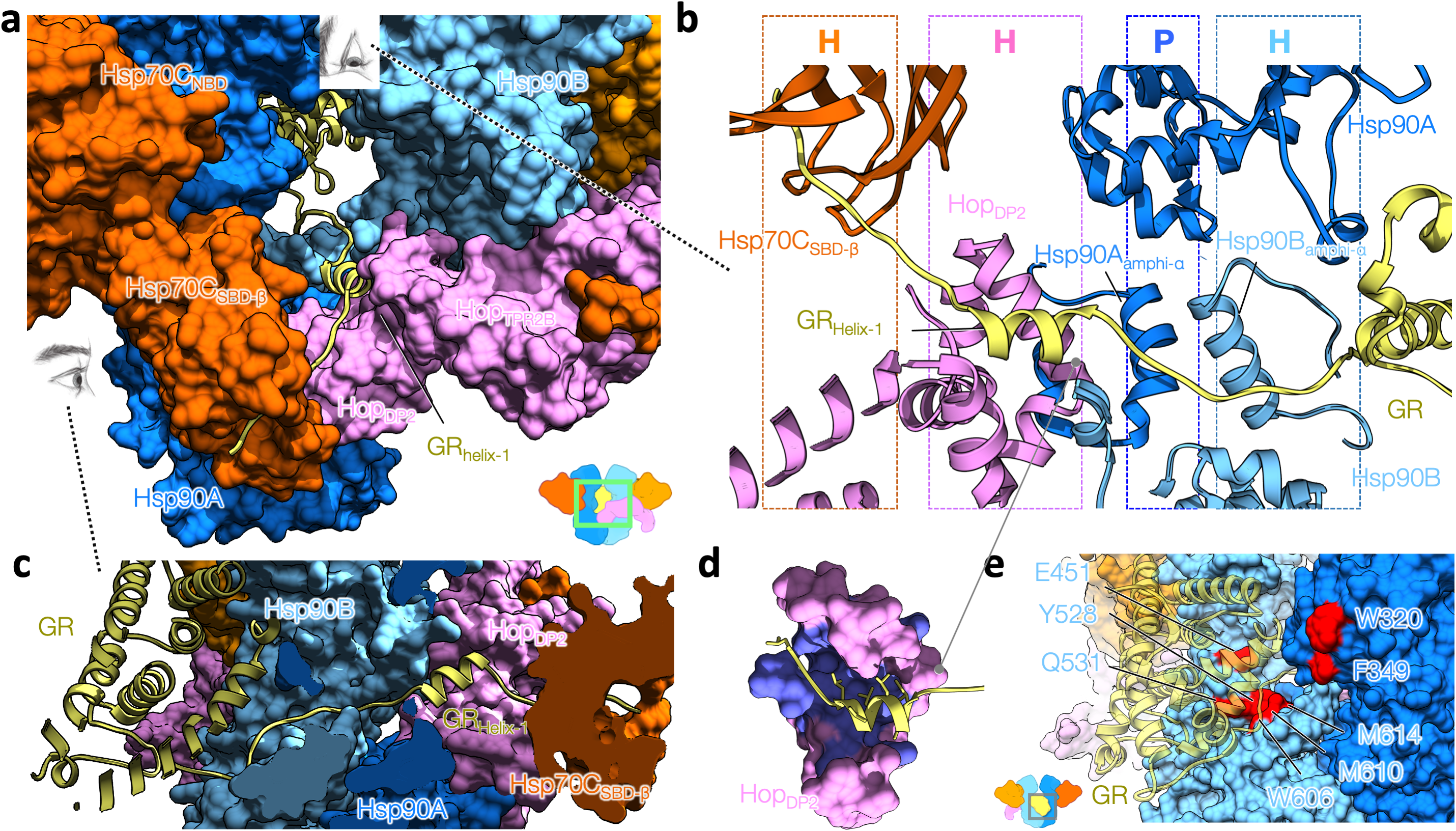
Facilitated by Hop, GR is loaded onto Hsp90 by Hsp70. **a,** Close-up front view of the loading complex (shown in surface representation). GR, shown in the ribbon model, is partially unfolded, with the N-terminal residues simultaneously griped by Hsp70C and Hop_DP2_, and is threaded through the semi-closed lumen of Hsp90. The remaining GR is at the other side of the loading complex; the ribbon model shown for the major body of GR is a modeling result from docking the GR crystal structure to the low-pass-filtered GR density. **b,** Top-to-bottom view of GR recognition via an extended client-binding pocket collectively formed by Hsp70C_SBD-β_, Hop_DP2_, and Hsp90A/B_amphi-α_s in ribbon representation. The N-terminal residues of GR (residues 517–533), which form a strand-helix-strand motif (yellow), are captured in the loading complex. The molecular properties provided by the individual binding pockets are color coded and labelled on the top panel (H and P denote hydrophobic and polar interactions, respectively). **c,** Side view of the GR N-terminal motif captured by the loading complex. **d,** Hop_DP2_, shown in surface representation, binds the LXXLL motif of GR Helix1, in which the hydrophobic residues of Hop_DP2_ are colored with purple and those of GR are shown in sticks. **e**, Residues on Hsp90 (surface representation) previously reported to be important for GR (transparent yellow ribbon) activation are highlighted in red.

Despite extensive 3D classifications, the main body of GR remained at low resolution. Nonetheless, the Hsp90_MD_s from each protomer and the Hsp90B_amphi-α_ clearly contact GR (Extended Data Fig. 22a,b,c,d). Hsp90 residues previously found^34,36,45,46,67^ to impact GR maturation are highlighted in Fig. 4e. The exposed Hsp90A^W320,F349^ directly contacts GR in both the loading complex (Fig. 4e, Extended Data Fig. 23c), and the maturation complex (Noddings et al., 2020). Notably, Hsp90^W320 (W300 in yHsp82)^ is an important binding residue exploited by both clients and cochaperones. Not only does it interact with GR and Hop_DP2_ (Fig. 3d), but also with another cochaperone Aha1^68^. Supporting its broad functional importance, numerous studies have reported effects of Hsp90A^W320^ mutations on GR activation^34,49,69^.

## Discussion

Our client-loading complex structure provides the first view of how Hsp70, Hsp90 and Hop work together to chaperone a client. Several features were unexpected: 1) two Hsp70s bind the Hsp90 dimer, one delivers client and the second scaffolds Hop. 2) Hop interacts extensively with all components, including GR, going well beyond the anticipated TPR-EEVD interactions. 3) Together Hop:Hsp90 and Hsp70:Hsp90 interactions define the Hsp90 conformation poising it for both client binding, and ultimately for ATP hydrolysis and client activation. 4) Hsp90 repurposes one side of its client-binding sites to bind Hop_DP2_, which in turn augments the Hsp90 lumenal clientbinding site, facilitating client-loading from Hsp70 (Fig. 5a).

**Fig. 5.**
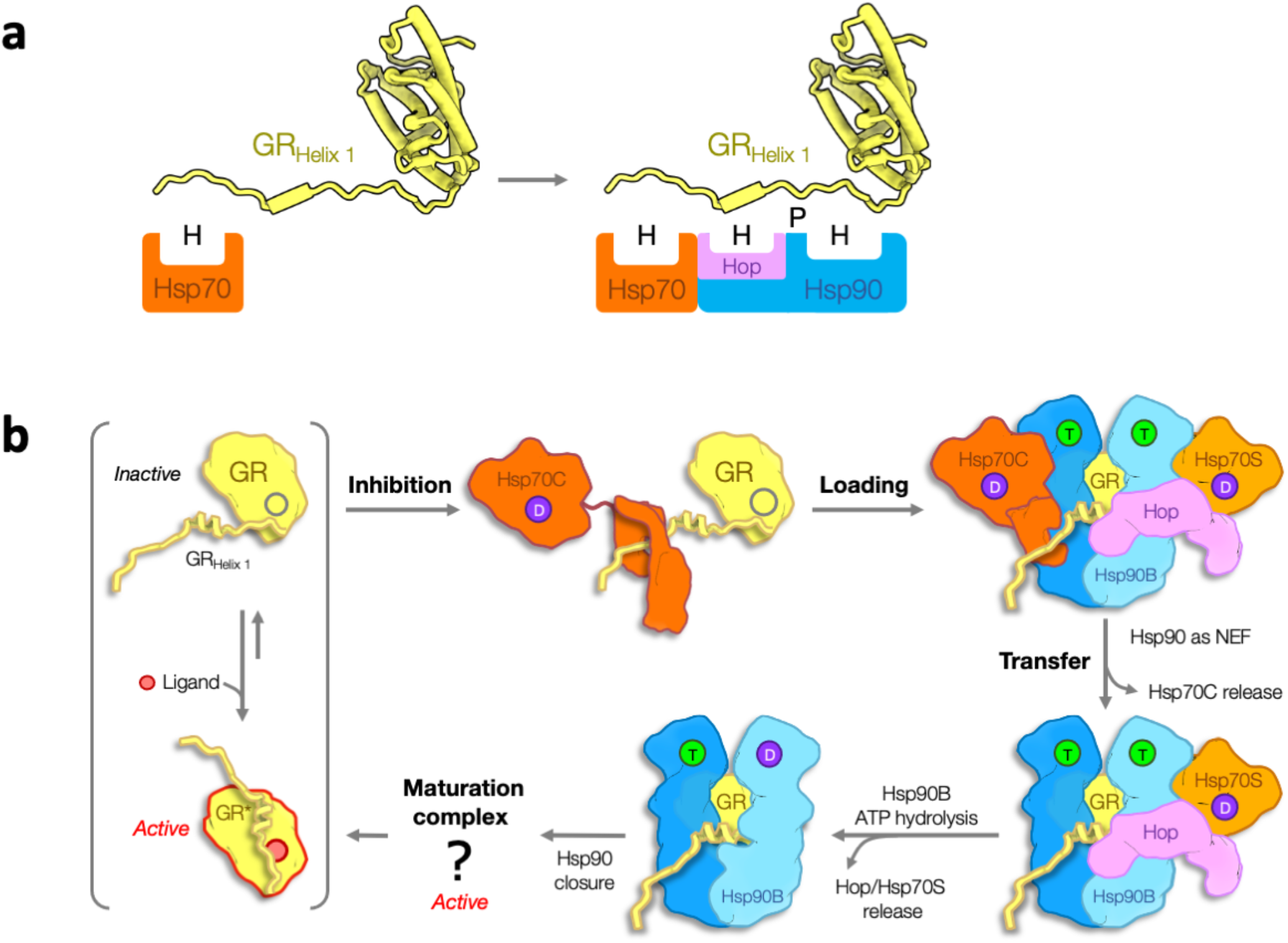
Schematic model of GR loading onto Hsp90 by Hsp70. **a,** The molecular principle of GR recognition at the client-loading step. Left, Hsp70_SBD_ (orange) inhibits GR (yellow), providing mostly hydrophobic binding (H). Right, the client-loading complex further stabilizes GR via an extended binding pocket assembled by Hsp70_SBD_ (orange), Hop_DP2_ (pink) and the lumen of Hsp90 dimer (blue), where stronger and more versatile molecular recognitions are provided for both hydrophobic (H) and polar (P) interactions. **b,** Molecular mechanism of GR transfer from Hsp70 to Hsp90 and the use of ATP hydrolysis on Hsp90. GR in physiological conditions is in an equilibrium of active and inactive states, in which the Helix 1 motif acts as a lid to stabilize ligand binding when attached. Hsp70C (dark orange) in its ADP state (D) binds GR’s pre-Helix 1 strand, facilitating the following motif to detach and hence inhibit GR (top-middle). Facilitated by Hop_DP2_ recognizing the LXXLL motif on GR Helix 1, Hsp70C loads GR to Hsp90 (blue), forming the client-loading complex (top-right). Although the structure was determined in an Hsp90^Apo^ state, we reason that in physiological conditions the high abundance of ATP would soon occupy Hsp90’s ATP binding pockets (T). Hsp90’s ATP binding and NEF activity facilitate Hsp70C release (bottom-right). The energy from the ATP hydrolysis (D) on Hsp90B (light blue) is used to release Hsp70S (light orange) and Hop (bottom-middle), followed by full closure of Hsp90.

The loading complex provides an extraordinarily extended client-binding pocket, with a large and very adaptable surface for client recognition (Fig. 4b and Extended Data Fig. 22d): (1) Hsp70 binds a hydrophobic strand, (2) Hop_DP2_ binds a hydrophobic/amphipathic helix, (3) the remaining part of the Hsp90A_amphi-α_ provides polar interactions, (4) the Hsp90B_amphi-α_ provides a hydrophobic surface, and (5) the Hsp90A/B lumen provides a combination of hydrophobic and polar interactions. Not only is the loading complex lumen spacious enough to bind a strand (as shown here) or intact helix (Extended Data Fig. 28), but the flexible positioning of the Hsp70C_SBD_ and the dynamic, adaptable conformation of the Hsp90_amphi-α_ allow even broader flexibility for client recognition (Fig. 4b,5a).

Which GR segment is captured in the loading complex lumen? The GR maturation complex structure unambiguously demonstrates that GR’s pre-Helix 1 region (GR^517–533^, GR_pre-Helix 1_) is gripped in the closed Hsp90 lumen (Noddings et al., 2020).

Reexamination of previous Hsp70-GR HDX-MS data^14^ reveals that only GR_pre-Helix 1_ becomes protected upon Hsp70 binding (Extended Data Fig. 24b) and GR_pre-Helix 1_ also contains high-scoring predicted Hsp70 binding sites (GR^518–524^, Extended Data Fig. 24a), strongly supporting that GR_pre-Helix 1_ is captured by Hsp70 in the loading complex (Extended Data Fig. 24b,c).. Moreover, a previous optical-tweezer study^70^ demonstrated that GR_Helix 1_ is readily detached, correlating with ligand binding loss. Together, this implies that perturbing GR_Helix 1_ by Hsp70 or the loading complex leads to loss of GR ligand binding. Altogether, we propose the following pathway for loading complex formation (Fig. 5b): Hsp70C captures the flexible GR_pre-Helix 1_, causing the following dynamic helix-strand motif to detach thereby destabilizing the GR ligand-binding pocket. Hsp70C then delivers the partially unfolded GR to Hop:Hsp70S:Hsp90. In the resultant loading complex, GR is further unfolded via engagement of GR_Helix 1_’s LXXLL motif with Hop_DP2_ and the GR_post-Helix 1_ strand with the Hsp90 lumen (Fig. 4b,d and Extended Data Fig. 27), suppressing any possible ligand binding. The rest of GR remains globular and is only loosely associated with the distal surface of Hsp90.

How does the loading complex progress to the maturation complex—a process requiring Hsp90 ATP hydrolysis and release of Hop and both Hsp70s? The one-Hsp70 loading complex suggests that the process is asymmetric and sequential, with the loss of Hsp70C occurring first, while Hsp90 ATP hydrolysis drives the release of the more tightly engaged Hsp70S-Hop. Schematically shown in Fig. 5b, we propose that a combination of Hsp90’s ATP binding and NEF activities promotes Hsp70C to release GR and exit the complex. This leaves GR engaged with Hop_DP2_ and the Hsp90 lumen, minimizing reformation of an Hsp70:GR complex or premature release. Lastly, as discussed in detail in Noddings et al., the conversion of the semi-closed Hsp90 in the loading complex to the fully closed Hsp90^ATP^ in the maturation complex may serve as a driving force for client remodeling and hence activation.

Contrary to kinases utilizing a dedicated cochaperone (Cdc37), GR uses a generalized chaperone (Hsp70) and cochaperone (Hop) for loading Hsp90 clients, making the principles learned here broadly applicable to other clients. Although Hop is absent in bacteria and organellar compartments, Hsp70s are present and the client-binding provided by Hop_DP2_ is likely substituted by the Hsp90_amphi-α_. While most, if not all, proteins engage with Hsp70 at least during initial folding, only a subset are Hsp90 clients. Ultimately, client properties must dictate this selectivity. Rather than an overall client property such as stability, our loading complex structure suggests a more nuanced balance of three effects: 1) the probability of partial unfolding fluctuations in the client, 2) the ability of Hsp70 to capture a transiently exposed site, and 3) the likelihood that further unfolding events would uncover adjacent client regions that can be captured by Hop_DP2_/Hsp90. Experiments to test these general principles can now be designed to predict and identify potential Hsp90/Hsp70 clients.

## Materials and Methods

### Protein purification

All recombinant chaperone proteins of Hsp90α, Hop, and Hsp70 (from human), and ydj1 (yeast Hsp40) were in general expressed and purified as described previously^14^ but with minor modifications as described below. Proteins were expressed in *E coli* BL21 star (DE3) strain. Cells were grown in TB at 37°C until OD_600_ reached 0.8. Protein expression was induced with 0.5 mM IPTG for 16 hours at 16°C. Cells were harvested by centrifugation at 4000xg for 15 minutes and resuspended in lysis buffer (50 mM Tris pH 7.5, 500 mM KCl, 10 mM Imidazole, 5mM βME). A protease inhibitor cocktail (Roche) was then added. Cells were lysed by an Emulsiflex system (Avestin). Lysates were cleared by centrifugation at 20,000 rpm for 1 hour at 4°C and the soluble fraction was affinity purified by gravity column with Ni-NTA affinity resin (QIAGEN). The protein was eluted by 50 mM Tris pH 8, 50 mM KCl, and 5 mM βME. The 6x-His-tag was removed with TEV protease and dialysis into low-salt buffer overnight (50 mM Tris pH 8, 50 mM KCl, and 5 mM βME). The cleaved protein was purified with MonoQ 10/100 GL (GE Healthcare), an ion-exchange column with 30 mM Tris pH 8, 50 mM KCl, 5 mM βME and eluted with a linear gradient of 50-500 mM KCl. Fractions with the target protein were then pooled and concentrated for final purification of size exclusion in 30 mM HEPES pH 7.5, 50 mM KCl, 2 mM DTT, and 10% Glycerol using a Superdex S200 16/60 (GE Healthcare) or Superdex S75 16/60 (GE Healthcare). The peak fractions were pooled, concentrated to ~100-150 μM or greater, snap-frozen in liquid nitrogen, and stored in aliquots at −80°C. MBP-GRLBD (F602S) was expressed and purified as described previously^14^. Note that for complex preparation, Hsp70 from Sf9 cell source was used with purification as described previously^14^. The Hop construct for the crosslinking experiment was obtained using quick-change at Q512 to the Amber codon. The construct was expressed in *E. coli* BL21 DE3 cells containing the pEVOL-pBpF plasmid^71^ distributed by the lab of Peter Schultz through Addgene (#31190). Cells were grown in terrific broth to an OD_600_ of 0.6. For induction, arabinose (0.02%), IPTG (1 mM), and *p*-benzoylphenylalanine (pBpa; 0.7 mM) was added, and expression was carried out overnight at 16°C. Cell harvesting, lysis, and Ni-NTA purification was performed as described above.

### Complex preparation

Using reaction buffer containing 50 mM HEPES pH 7.5, 50 mM KCl, and 2 mM DTT, 10 μM Hsp90 dimer of D93A mutant, 10 μM Hop, 15 μM Hsp70, 4 μM Hsp40, and 20 μM MBP-GRLBD were incubated with 5mM ATP/MgCl for 1 hour at room temperature. The complex was purified and analyzed by SEC-MALS with a Wyatt 050S5 column on an Ettan LC (GE Healthcare) in a running buffer containing 50 mM HEPES, 50 mM KCl, 5 mM MgCl2, 2 mM DTT, 200 μM ADP and 0.01% Octyl β-D-glucopyranoside (β-OG); Molecular weights were determined by multiangle laser light scattering using an in-line DAWN HELEOS and Optilab rEX differential refractive index detector (Wyatt Technology Corporation). Once eluted, fractions containing the GR-loading complex were immediately crosslinked with 0.02% glutaraldehyde for 20 minutes at room temperature and quenched with 20 mM Tris pH 7.5. Fractions containing the GR-loading complex were separately snap-frozen in liquid nitrogen, and stored in aliquots at −80°C.

### Photoreactive crosslinking experiment

To ensure that Hop crosslinks the bound segment in the context of the loading complex, crosslinking reactions were performed immediately after the complex was fractionated from SEC (see the above complex preparation section). Using a UV-transparent, 96- well microplate (Corning) as a fraction collector, the whole fractions of the eluted GR-loading complex were subjected to UV exposure using an agarose gel imaging system (Enduro GDS Imaging System). Samples were irradiated for 60 mins in total. To prevent overheating, the 96-well plate was placed on a shallow plate filled with constantly refreshed ice water during the time course of the exposure. SDS-PAGE was used to analyze cross-linked product, followed by Western plot transfer to nitrocellulose and probed with an MBP antibody (New England BioLabs) (Extended Data Fig. 26).

### Cryo-EM sample/grid preparation and data acquisition

The flash-frozen fractions of the loading complex were thawed and concentrated to 0.7-0.8 μM. About 2.5 uL of the complex sample was applied onto a glow-discharged, holey carbon grid (Quantifoil R1.2/1.3, Cu, 400 mesh), blotted by Vitrobot Mark IV (FEI) for 8-14 seconds at 10°C / 100% humidity, and plunge-frozen in liquid ethane. Four data collections were made using Titan Krios (Thermo Fisher Scientific) equipped with K2 camera (Gatan K2). SerialEM^72^ was used for all the data collections with parameters as described in Extended Data Table 1.

### Image processing

Movies were motion-corrected using MotionCor2^73^, in which the unweighted summed images were used for CTF estimation using CTFFIND4^74^, and the dose-weighted images were used for image analysis with RELION^42^ throughout. The initial model of the loading complex was obtained from a small data collection (Extended Data Fig. 2a). Particles were picked from the small data collection using Gautomatch (https://www2.mrc-lmb.cam.ac.uk/research/locally-developed-software/zhang-software/) without template and subjected to reference-free 2D classification *(Class2d)* using RELION. 2D class averages with proteinaceous features were selected for 3D classification *(Class3d)* using RELION. For *Class3d,* the reference model was generated using the semi-open conformation Hsp90 from the Hsp90:Hop cryoEM structure^63^ (Extended Data Fig. 21). Among eight classes, one class (~8Å resolution) showed recognizable shapes of the protein components, although the class is drastically different from the initial model. This low-resolution reconstruction of the loading complex was then used as an initial reference model for the following image analysis that achieved high-resolution.

The procedure to obtain the high-resolution reconstruction is shown schematically in Extended Data Fig. 2b. Particles were picked from all dose-weighted micrographs using Gautomatch with the low-resolution reconstruction as a template. Without using *Class2d,* the extracted, binned 4 × 4 particles (4.236 Å pixel^-1^) were subjected to RELION *Class3d* (4 classes) to sort out “empty” or non-proteinaceous particles. Particles from the selected class were re-centered and re-extracted to binned 2 × 2 (2.118 Å pixel^-1^) for another round of *Class3d.* Note that a low-resolution (~7.5Å) reconstruction of a one-Hsp70 loading complex was obtained among the 4 classes. The selected class that contains 636,056 particles of the two-Hsp70 loading complex was 3D auto-refined *(Refine3d)* into a single class *(consensus class*). The set of particles are then used for further global classification and focused classification (described below). For global classification, another round of masked *Class3d* (4 classes) was performed without alignment, followed by masked *Refine3d* using unbinned particles (1.059Å pixel^-1^). Finally, 85,619 particles from the highest resolution, the two-Hsp70 loading class were further subjected to multiple rounds of per-particle CTF/beam-tilt refinement until the gold-standard resolution determined by *Refine3d* no longer improved. The overall gold-standard resolution for the global reconstruction of the loading complex is 3.57 Å (Extended Data Fig. 3). Local resolution was estimated using the RELION (Extended Data Fig. 3a).

The loading complex presents conformational heterogeneity at all regions of the complex, in particular at the Hop_TPR2A-TPR2B_ (Extended Data Fig 17. a-c). Starting from the consensus class containing 636,056 binned 2 × 2 particles (2.118Å pixel^-1^), masks at various GR-loading complex regions were used for focused classification with signal subtraction *(Focused-class3d;* Extended Data Fig. 4b). A pipeline to obtain the best reconstruction for each masked region is outlined in Extended Data Fig. 4a. For each masked region, *Class3d* without alignment was performed, followed by *Refine3d* using unbinned particles (1.059Å pixel^-1^). For each *Class3d* job, parameters of number of requested classes (*K*=6,8,10,12,14) and *Tau* (*T*=10,20,30,40) were scanned. For each masked region, the reconstruction that results in the highest resolution determined by the gold-standard FSC of the *Refine3d* job was selected for each masked region. Using the unbinned particles, another round of *Focused-class3d* was performed with the similar procedure described for the previous round. Parameters were scanned in a similar manner but with smaller requested classes (*K*=2,3,4,5). Finally, the selected *Focused-class3d* job was subjected to multiple rounds of per-particle CTF/beam-tilt refinement. The overall resolution of the reconstruction for each masked region is determined by the gold-standard FSC and as denoted in the FSC plots in Extended Data Fig. 4b. The focused maps showed much better atomic details than the global reconstruction at all regions, and hence were used for model building and refinement.

### Model Building and refinement

Model building and refinement was carried out using Rosetta throughout. All the components of the loading complex had crystal structures or close homologous structures (from yeast) available. Details of how the starting, unrefined atomic model for each component was obtained are described below. For Hsp90, the starting model was assembled from the crystal structure of human Apo-Hsp90NTD (PDB ID: 3T0H)^53^ and the cryo-EM structure of the Hsp90_MD-CTD_ from the GR-maturation complex (Noddings et al., 2020). Many crystal structures of Hsp70_NBD_ were available. As Potassium and Magnesium ions were used in the buffer for complex preparation and there is density accounted for them in our focused map (Extended Data Fig. 13), the ADP state Hsp70_NBD_ structure crystal structure (PDB ID: 3AY9)^54^ that has Potassium and Magnesium ions to coordinate ADP was used as an starting model. For the Hsp70_SBD_, the human Hsp70 crystal structure (PDB ID: 4PO2)^75^ was used as a starting model. For Hop, the crystal structures^61^ of the Hop_TPR2A-TPR2B_ (PDB ID: 3UQ3) and the Hop_DP2_ (PDB ID: 2LLW) from yeast were used as initial templates with the alignments obtained from HHpred server^76^ (Extended Data Fig. 16). The insertion in the threaded Hop model was completed using RosettaCM guided by the cryo-EM density^77^. The resulting completed models of Hop_TPR2A_ and Hop_TPR2B_ showed a high resemblance to their structures determined by NMR individually (Extended Data Fig. 16c,d). The sequence of the Hsp70C_SBD-β_-bound GR segment was determined with the aid of Rosetta (Extended Data Fig. 25). Two 7-residue GR segments (SIVPATL and IVPATLP) of a continuous sequence (residues 518-525; note that residue 518 in the native GR sequence is T, not S) in the pre-Helix 1 regions are predicted to be Hsp70 binding sites by two state-of-the-art algorithms (BiPPred^78^ and ChaperISM^79^). Structural modeling of the two GR peptides in the templates^80^ (PDB IDs: 4EZZ, 4EZT, and 4EZQ) with “reverse” binding mode of Hsp70C_SBD-β_ indicated that the segment, SIVPATL, is energetically more favored (Extended Data Fig. 25a-c).

Using the high-resolution information acquired from focused classification/refinement, the starting models were refined separately into the individual focused maps (Extended Data Fig 4.). Model overfitting was monitored and harbored using the half-map approach as previously described^81,82^, in which one-half map from RELION *Refine3d* was used for density-guided refinement while the other half map was used for validation. Rosetta fragment-based iterative refinement protocol^82^ was used to refine the models throughout. Based on the high-resolution focused maps, the refinement tasks were split into (1) Hsp90A:Hsp70C, (2) Hsp90B:Hsp70S (Extended Data Fig. 4,7), (3) Hsp70S:Hop_TPR2A_ (Extended Data Fig 4,18) and (4) Hsp90AB_CTD_:Hsp70S_SBD- β_:Hop_DP2_:GR_Helix 1_ (Extended Data Fig. 4,22). To model the GR segment threaded through the lumen of Hsp90, the GR_Helix 1_ motif (residues 528-551) was first segmented from the crystal structure of GR_LBD_ (PDB ID: 1M2Z)^83^ and rigid-body fitted into the lumen density. The GR_Helix 1_ segment was then rebuilt and refined using Rosetta fragment-based iterative refinement method into the focused map of the Hsp90AB_CTD_:Hsp70S_SBD-β_:Hop_DP2_:GR_Helix 1_ (Extended Data Fig. 22). The remaining, globular portion of GR_LBD_ was rigid-body fitted initially using Chimera. The placement was further refined, guided by (1) the connectivity to the GR_Helix 1_ motif and (2) GR’s interaction with Hsp90_MD_ in the maturation complex. The docked GR_LBD_ was then energy minimized in Rosetta guided by low-pass filtered cryo-EM map. Finally, the connectivities of the N-terminal end of the globular portion of GR_LBD_ and the C-terminal end of the GR_Helix 1_, and the N-terminal end of the GR_Helix 1_ motif and the C-terminal end of the Hsp70-bound GR_pre-Helix 1_ portion were built using RosettaCM. The final model of the loading complex was assembled and refined into the high-resolution global construction.

Structural modeling was used to ensure and suggest the connectivities of the EEVD tails of Hsp90/Hsp70 to the bound TPR domains of Hop (Extended Data Fig. 20). For each protomer of Hsp90, ~40 residues of the tail were modeled using RosettaCM to connect the bound Hsp90 EEVD fragment and the very C-terminal helix in the Hsp90CTD. Similarly, for Hsp70C, the remaining residues were built, including a Hsp70_SBD-α_ lid closing on the Hsp70_SBD-β_ (PDB ID: 4PO2)^75^ followed by ~30 residues tail residues to the Hsp70 EEVD fragment bound to Hop_TPR2B_.

### In vivo yeast Hsp90:Hsp70 interaction assay

Hsc82 plasmids expressing untagged or His-Hsc82 were expressed in yeast strain JJ816 *(hsc82::LEU2 hsp82::LEU2/YEp24-HSP82).* His-Hsc82 complexes were isolated as described ^84^. Antibodies against the last 56 amino acids of Ssa1/2 were a gift from Dr. Elizabeth Craig (University of Wisconsin). The Sti1 peptide antisera was raised amino acids 91-108. His-Hsc82 was detected using an anti-Xpress antibody, which recognizes sequences near the 6X-His tag at the amino-terminus. The plasmid pBv- src, which expresses v-src under the *GAL1* promoter, and the corresponding empty vector pB656 were a gift from Frank Boschelli ^85^. The R46G and K394E mutations were isolated in a genetic screen as described ^47^.

## Acknowledgements

We thank members of the Agard Lab for helpful discussions. We thank Dr. Tristan W. Owens for advising the photoreactive crosslinking experiment. We thank Michael Braunfeld, David Bulkley, Glenn Gilbert, Eric Tse, and Zanlin Yu from the W.M. Keck Foundation Advanced Microscopy Laboratory at the University of California San Francisco (UCSF) for maintaining the EM facility and help with data collection. We thank Matt Harrington and Joshua Baker-LePain for computational support with the UCSF Wynton cluster. R.Y.-R.W. was a Howard Hughes Medical Institute Fellow of the Life Sciences Research Foundation. The work was supported by funding from Howard Hughes Medical Institute (D.A.A.) and NIH grants R35GM118099 (D.A.A.), S10OD020054 (D.A.A.), S10OD021741 (D.A.A.), and R01GM127675 (J.L.J.).

## Author contributions

R.Y.-R.W. performed the research and drafted the manuscript. D.A.A. supervised the research. E.K. trained R.Y.-R.W. for the biochemistry of the GR reconstitution system. A.G.M. trained R.Y.-R.W. for cryo-EM operation and data acquisition. J.L.J. carried out *in vivo* yeast experiments. R.Y.-R.W., C.M.N., and D.A.A. wrote the manuscript with input from all authors.

---

**Extended Data Fig. 1.**
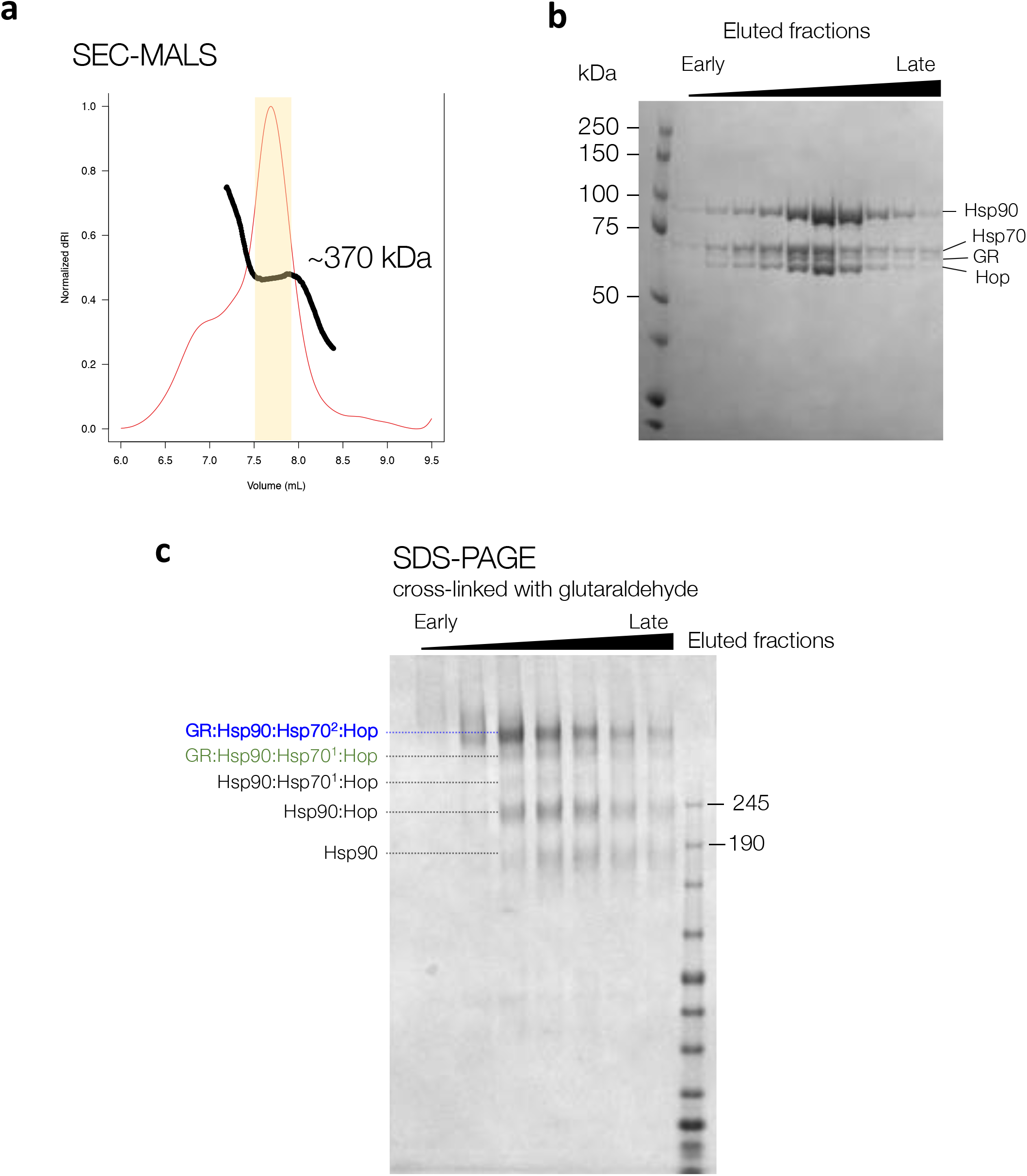
In vitro reconstitution and purification of the GR-loading complex. **a**, Elution profile of gel filtration using SEC-MALS to confirm the homogeneity of the GR-loading complex. The apparent molecular weight of the eluent estimated by SEC-MALS is ~370 kDa although the two-Hsp70 client-loading complex is ~440 kDa. The discrepancy may be a result of multiple species co-eluted. **c**, SDS-PAGE stained with Coomassie blue of the eluted fractions marked in (**a**). **d**, SDS-PAGE of the fractions treated with 0.02% (w/v) glutaraldehyde crosslinking for 20 minutes at room temperature, followed by quenching with 20 mM Tris buffer at pH 7.5.

**Extended Data Fig. 2.**
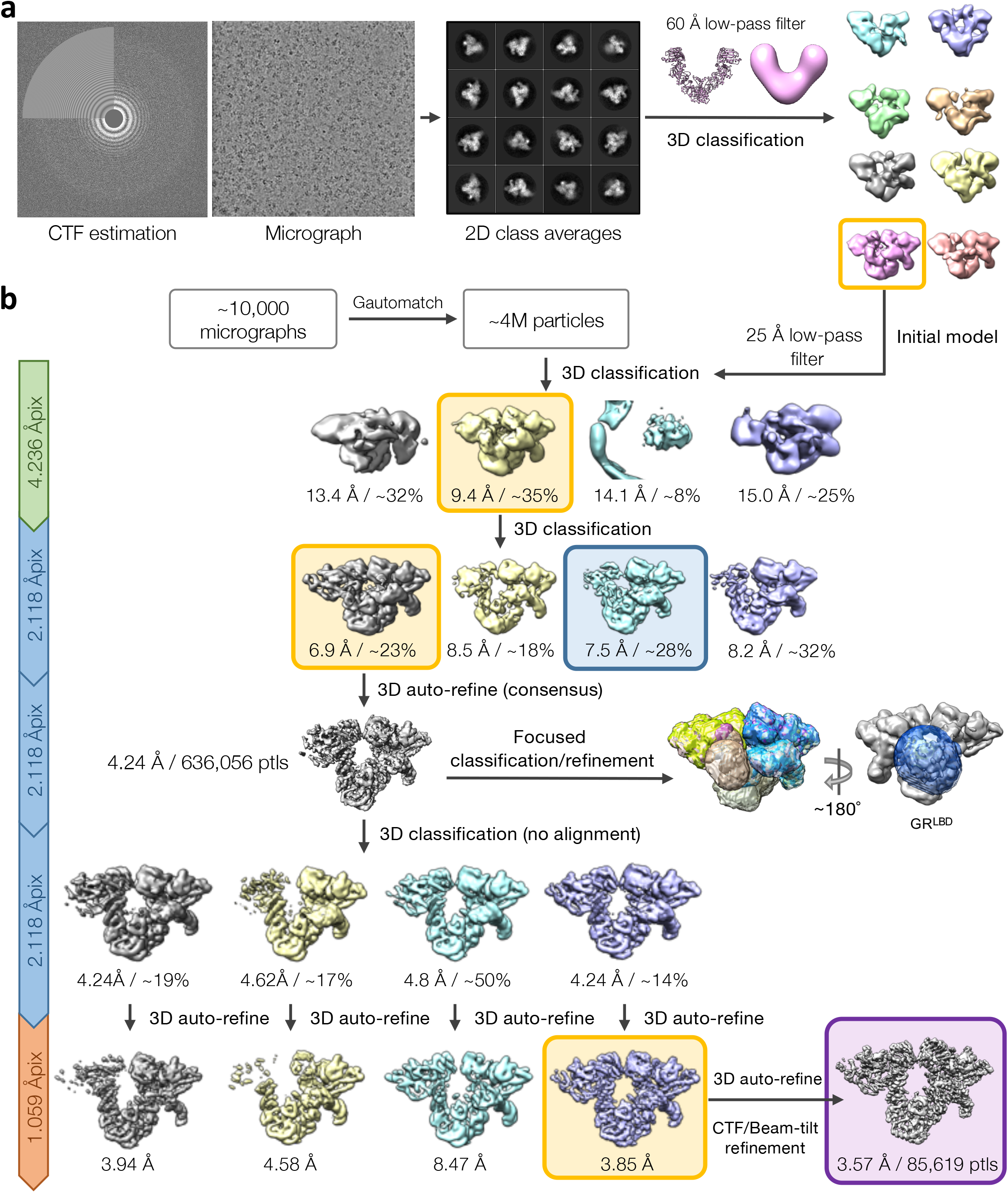
Cryo-EM single-particle image processing pipeline of the GR-loading complex. **a**, Initial model generation for the GR-loading complex. The 60 Å low-pass filtered initial model used to reconstruct the 3D model was adopted from the Hsp90 semi-open conformation structure from the Hsp90:Hop cryo-EM complex (Southworth & Agard, 2011). **b**, Schematic workflow of the global cryo-EM map reconstruction. Yellow boxes indicate the selected class to move forward. Blue box indicates one-Hsp70 loading complex. Purple box indicates the final high-resolution global reconstruction.

**Extended Data Fig. 3.**
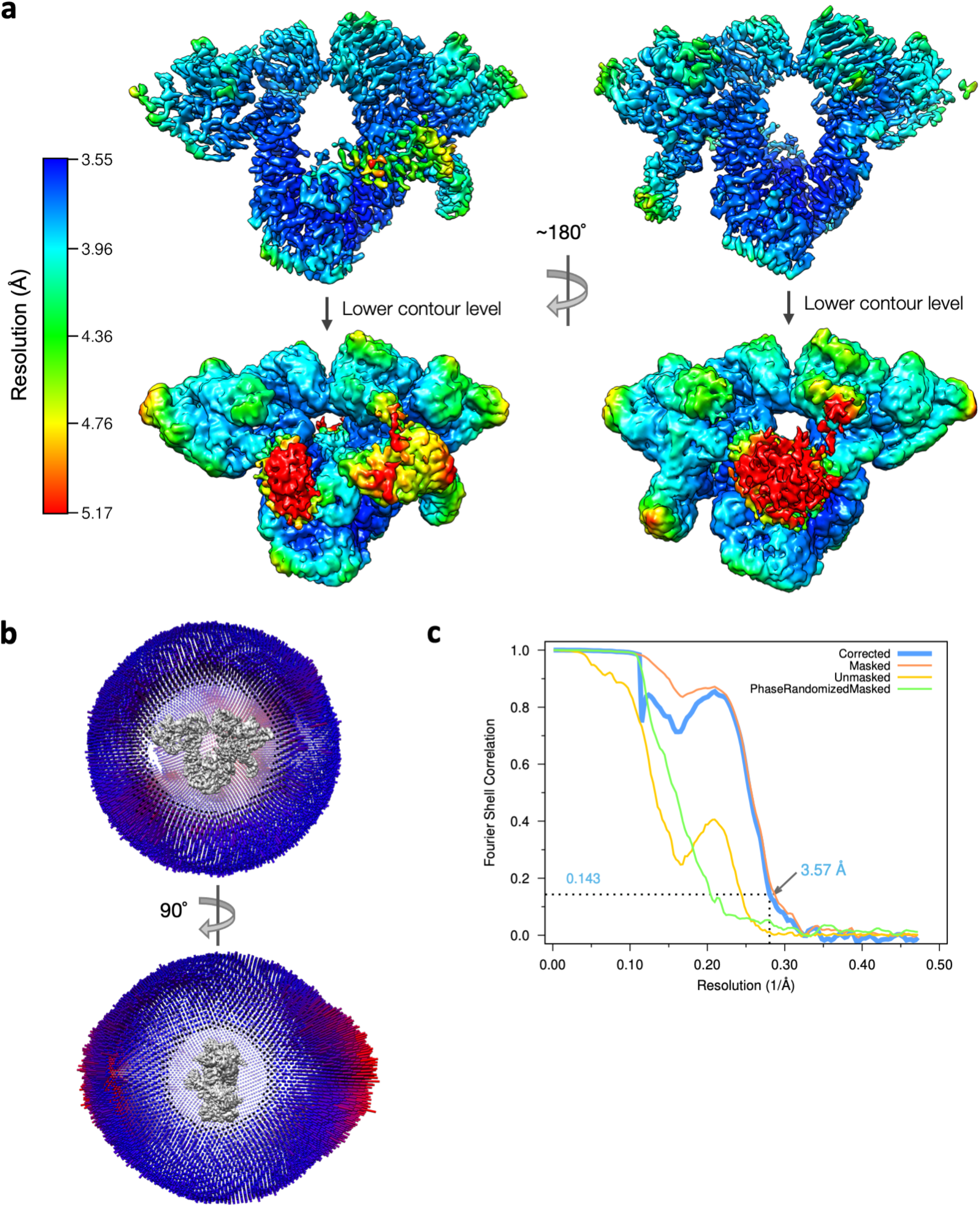
Cryo-EM single-particle analysis of the GR-loading complex. **a,** Local resolution estimates for the GR-loading complex global reconstruction were calculated using RELION with front view (left) and back view (right). **b,** Euler angle distribution in the final reconstruction. Orthogonal views of the reconstruction are shown with front view (top) and side view (bottom) **c,** Gold Standard FSC for the global cryo-EM reconstruction (top).

**Extended Data Fig. 4.**
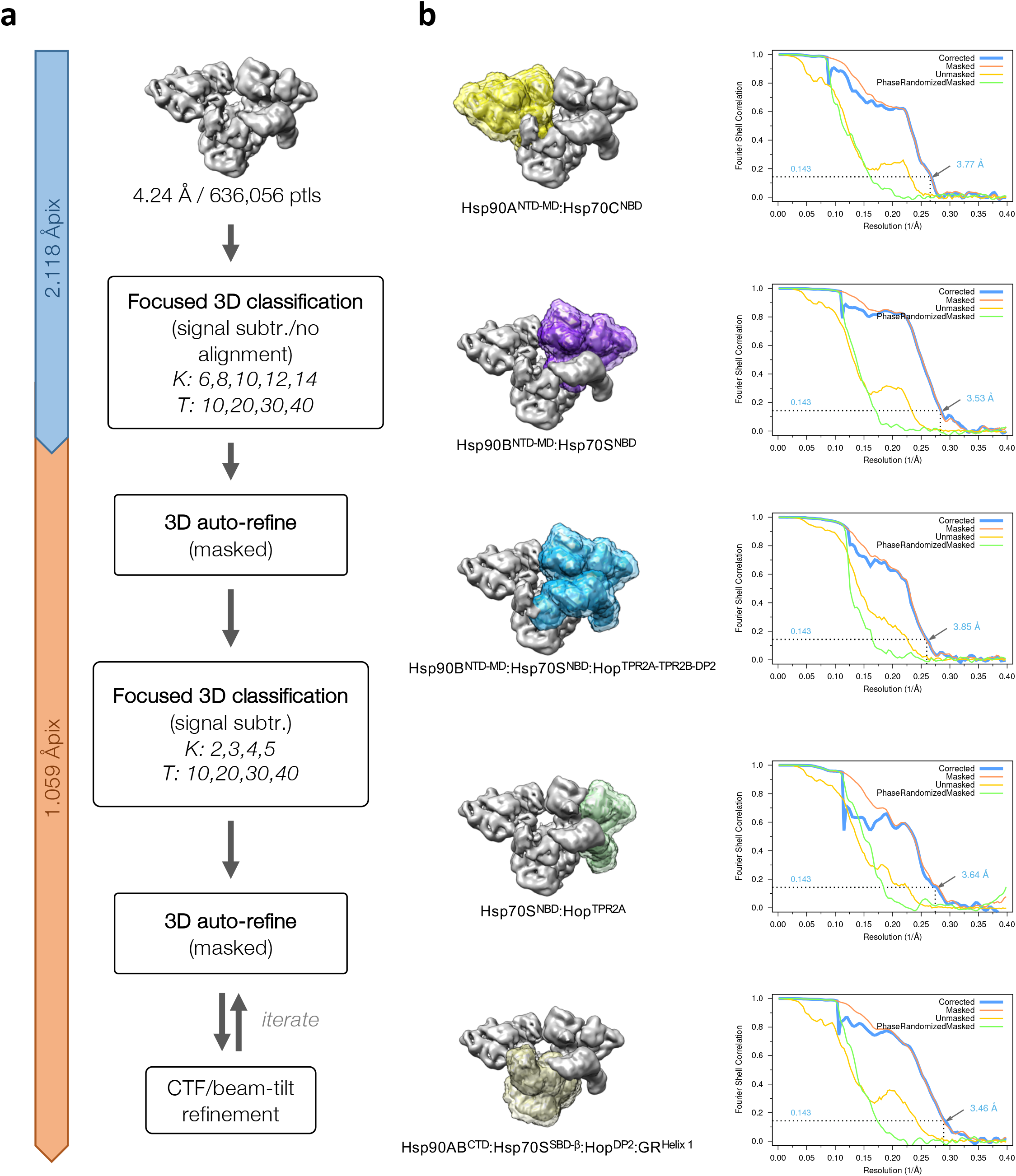
Focused classification and refinement. **a**, Flow chart of focused classification/refinement using the signal subtraction approach. Final reconstructions for individual masked classifications/refinements were selected based on the resolution intercepted with the FSC 0.143 from 3D auto-refine. **b**, Masks were created at various regions of the GR-loading complex (left) and its corresponding Gold Standard FSC (right) after 3D auto-refine. The nominal resolution for each reconstruction is labeled and indicated in the FSC plots.

**Extended Data Fig. 5.**
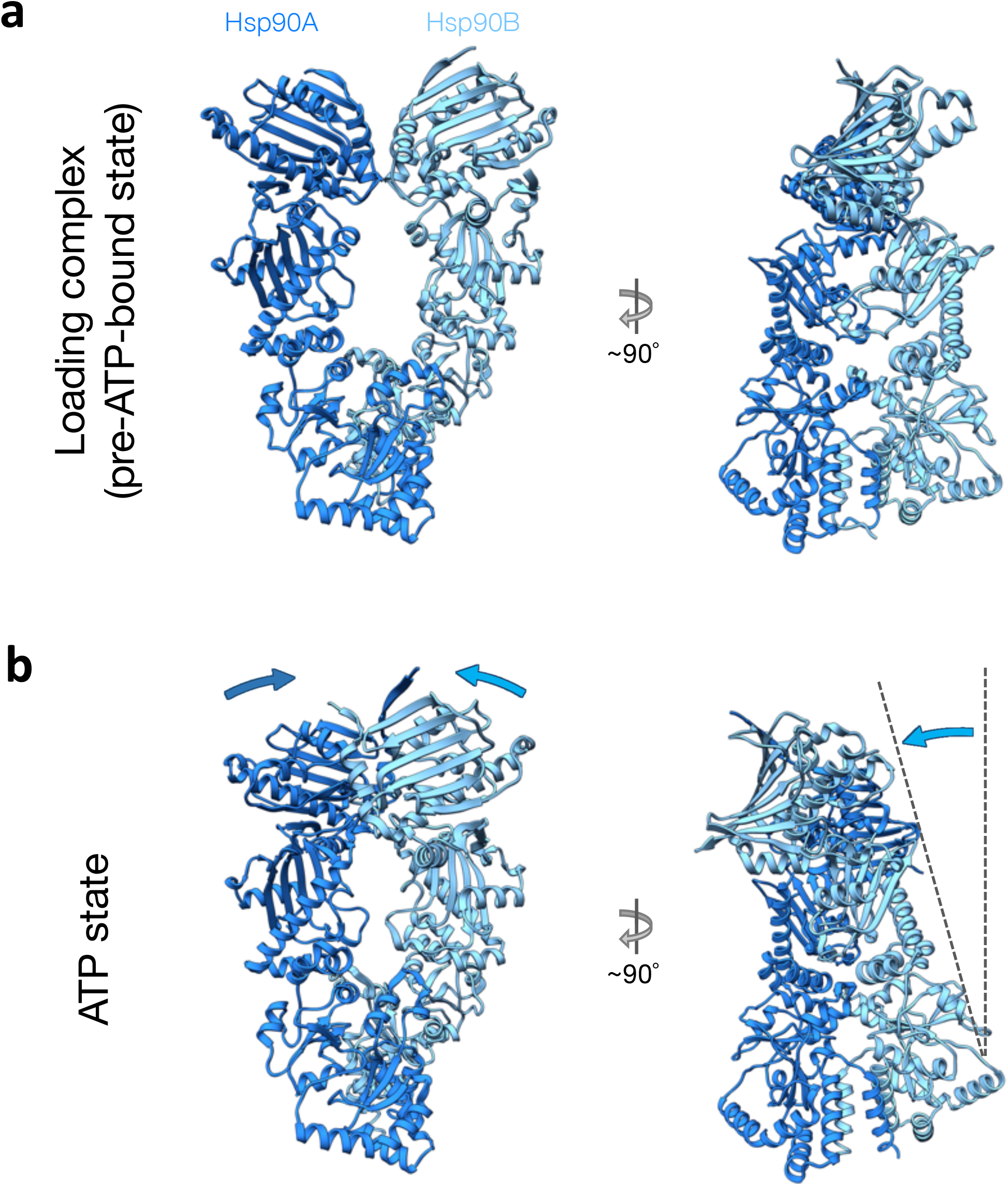
The Hsp90 in the loading complex is one step away from the fully closed ATP state. **a**, Front (left) and side (right) views of the Hsp90 in the loading complex. **b**, Front and side views of Hsp90 in the ATP state (the Hsp90 in the GR-maturation complex). Arrows indicate displacements from the Hsp90 in the loading state, in which a large twisting motion is apparent from the side view.

**Extended Data Fig. 6.**
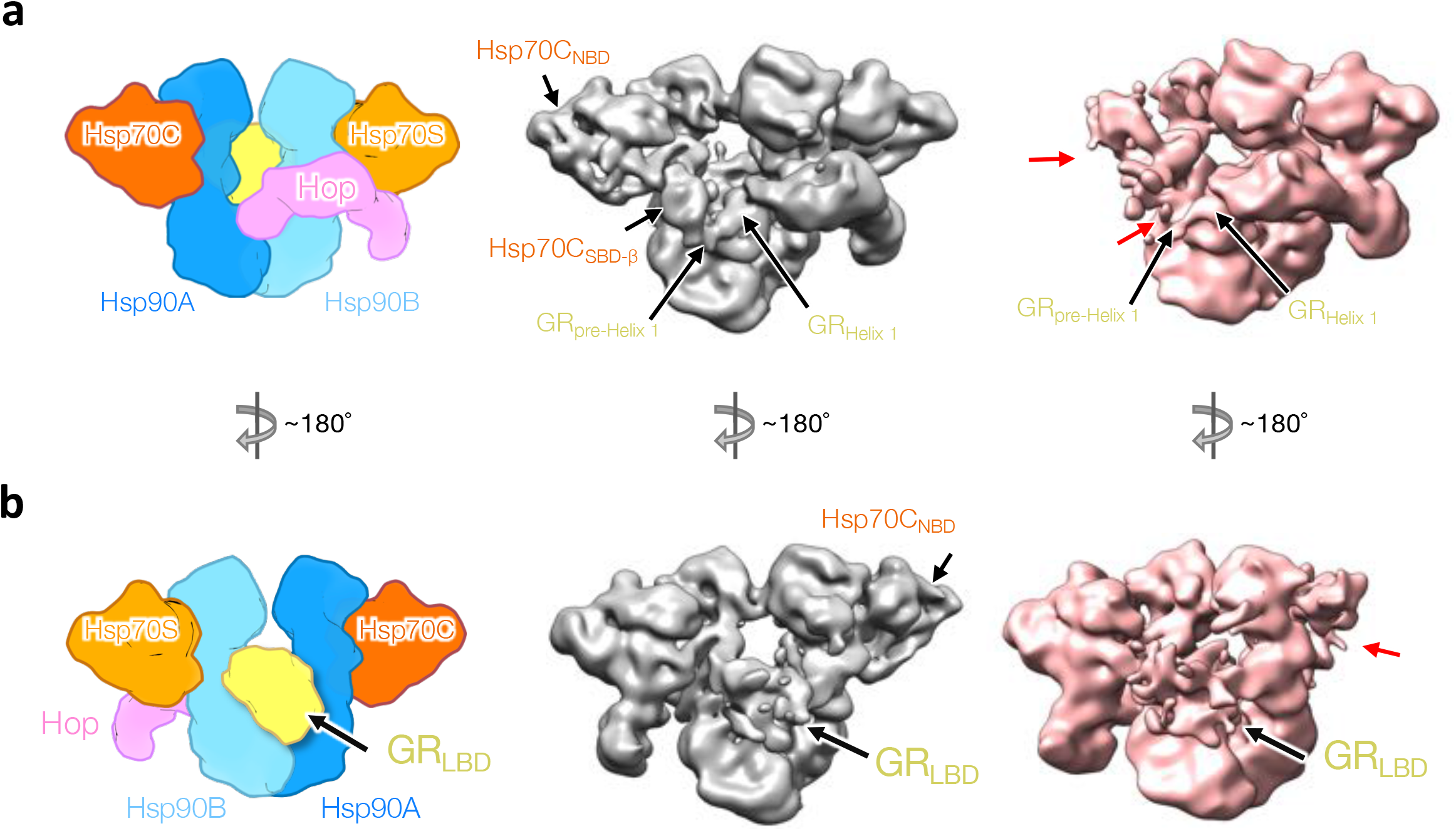
One-Hsp70 GR-loading complex. **a**, Schematic model of the two-Hsp70 loading complex (left). Front views of cryo-EM maps of the two-Hsp70 (middle; gray color) and one-Hsp70 (right; salmon color) GR-loading complexes. Right, the one-Hsp70 reconstruction has lost density for Hsp70C NBD and Hsp70C SBD-α (red arrows); however, density for GR_pre-Helix 1_ and GR_Helix 1_ is in the same locations as it is in the two-Hsp70 GR-loading complex. **b**, 180 degree views from the top.

**Extended Data Fig. 7.**
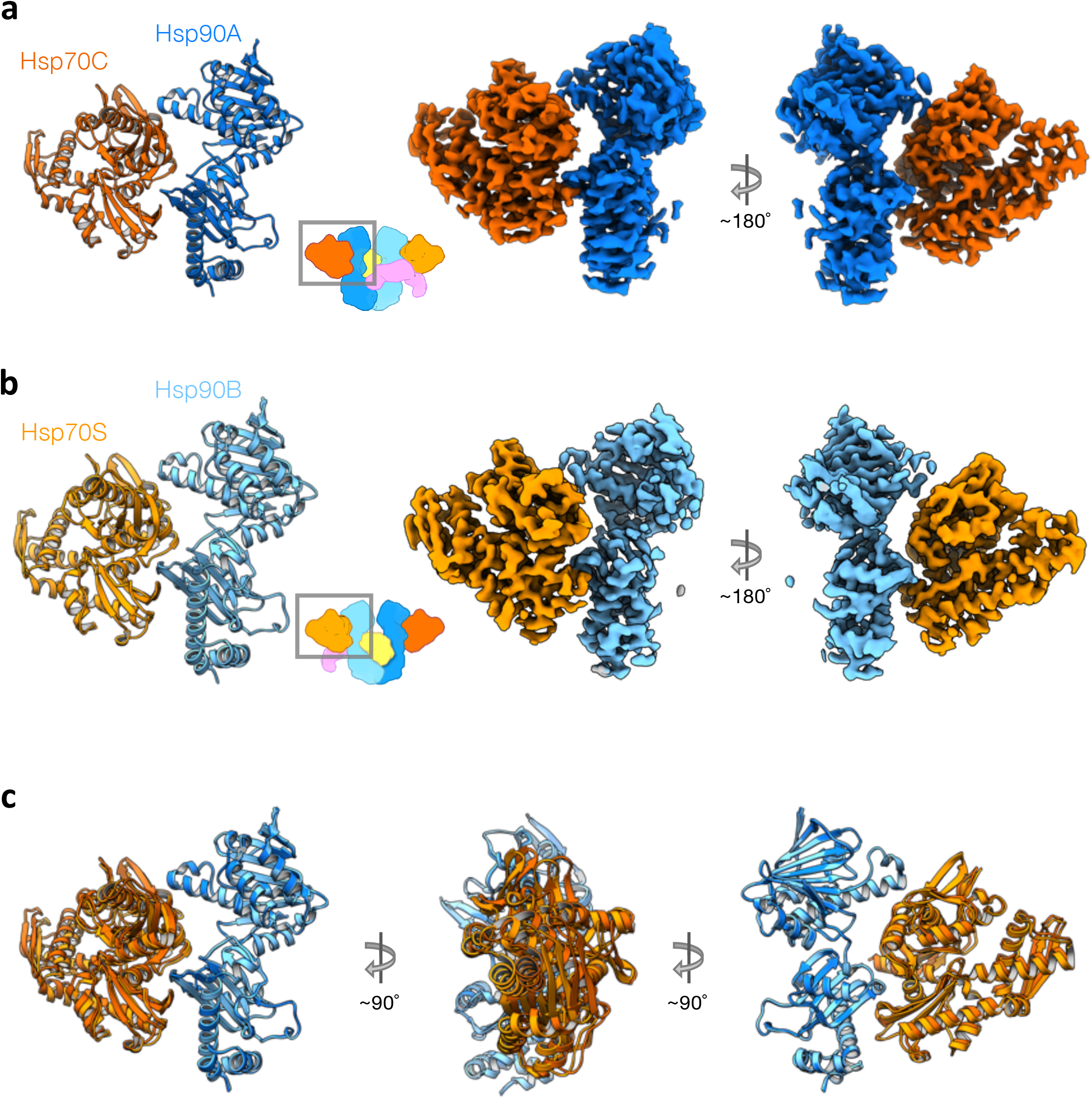
Hsp90 and Hsp70 pairs in the GR-loading complex. **a**, Hsp90A and Hsp70C are shown in ribbon representation (left) colored in dark orange and dark blue, respectively. The corresponding focused map (middle and right) with a 3.77Å resolution. **b**, Hsp70B and Hsp70S are shown in ribbon representation (left) colored in orange and blue, respectively. **c**, Overlay of the Hsp90:Hsp70 pairs from (**a**) and (**b**) with Hsp90s aligned.

**Extended Data Fig. 8.**
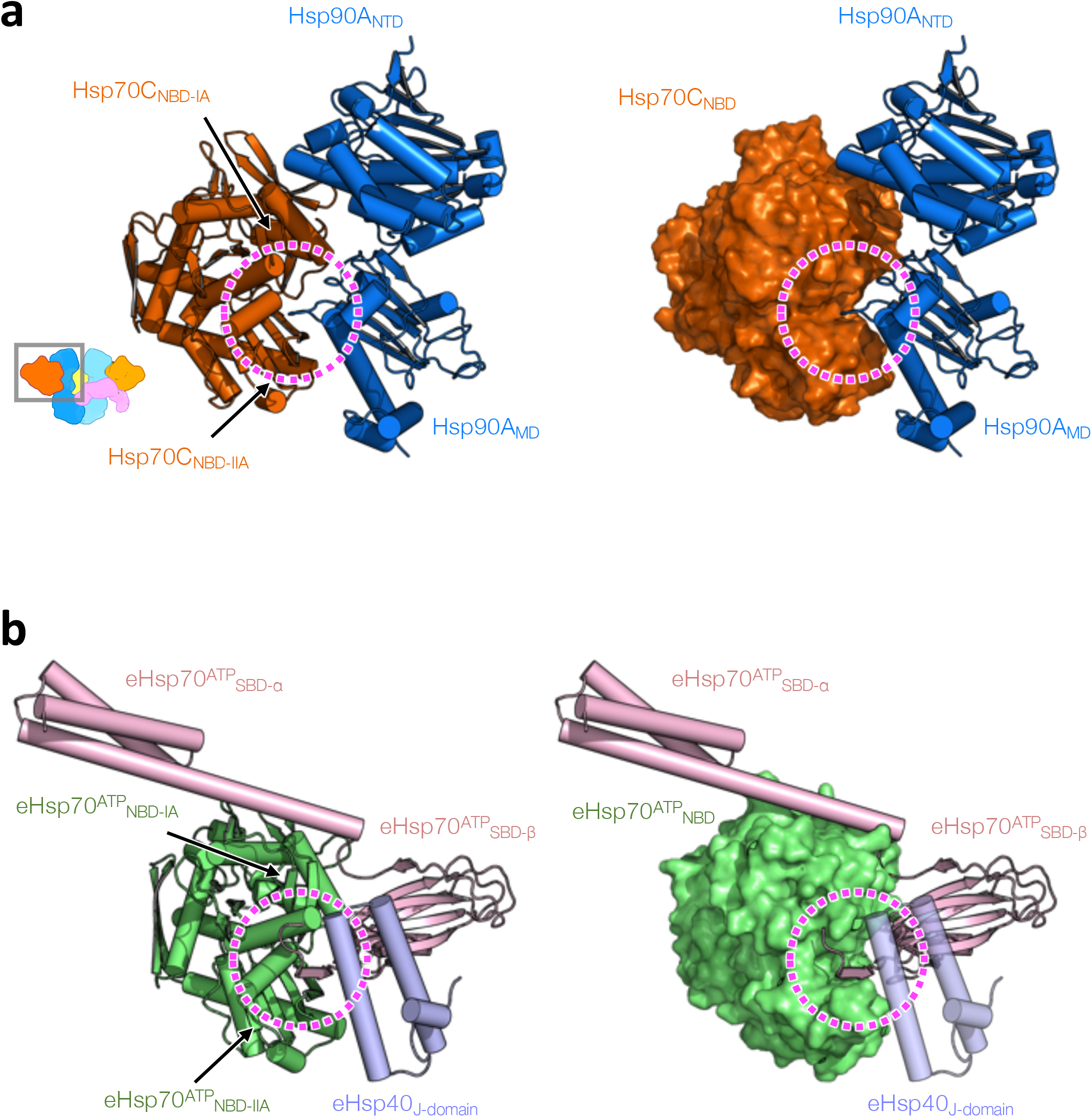
The Hsp70 cleft, formed by Hsp70_NBD-IA_ and Hsp70_NBD-IIA_, which Hsp90MD interacts with, is used by the interdomain linker in the Hsp70 ^ATP^ state and Hsp40’s J-domain. **a**, Cartoon (left) and surface (right) representation of Hsp90A:Hsp70C in the loading complex. The Hsp70 cleft is indicated by dashed circles. **b**, Cartoon (left) and surface (right) representation of the *E. coli* Hsp70 (eHsp70):J-protein complex in the ATP state (PDB ID: 5nro). The two subdomains of Hsp70 are colored in green for eHsp70^ATP^_NBD_ and in pink for eHsp70^ATP^_SBD_. The cleft that the interdomain link binds is indicated by dashed circles.

**Extended Data Fig. 9.**
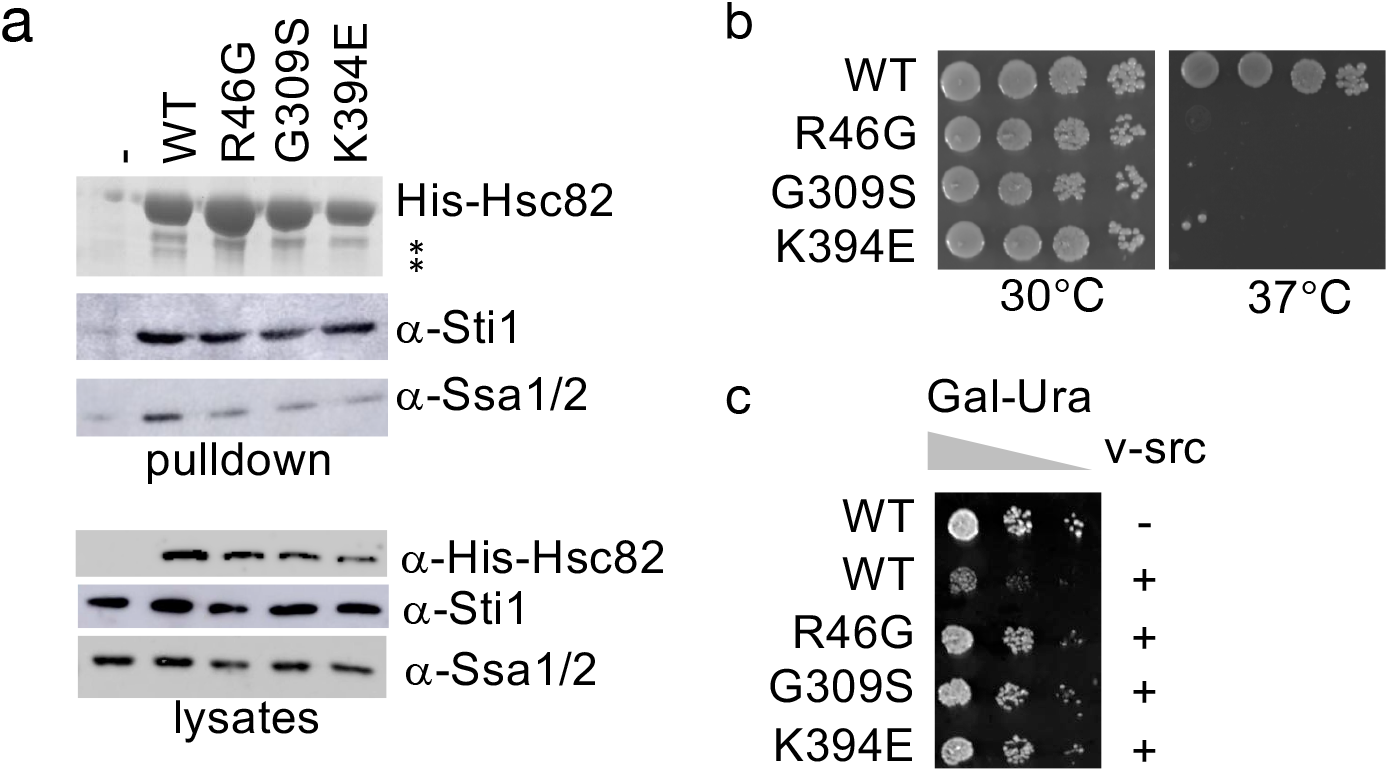
Mutations that disrupt Hsp90:Hsp70 interactions in yeast. **a**, His-Hsc82 complexes were isolated from yeast and analyzed by SDS-PAGE and visualized by Coomassie staining and immunoblot analysis. Yeast proteins: Sti1=Hop; Ssa1/2=Hsp70; Hsc82=Hsp90β. **b**, Plasmids expressing wild-type or mutant Hsc82 were expressed as the sole Hsp90 in JJ816 *(hsc82hsp82)* cells. Growth was examined by spotting 10-fold serial dilutions of yeast cultures on rich media, followed by incubation for two days at 30°C or 37°C. **c**, Strains expressing WT or mutant *HSC82* were transformed with a multicopy plasmid expressing *GAL1*-v-src (pBv-src) or the control plasmid (pB656) (Dey et al. 1996). Yeast cultures were grown overnight at 30° in raffinose-uracil drop-out media until mid-log phase. 20% galactose was added to a final concentration of 2%. After six hours, cultures were serially diluted 10-fold onto uracil drop out plates containing galactose. Plates were growth for 2-3 days at 30°C.

**Extended Data Fig. 10.**
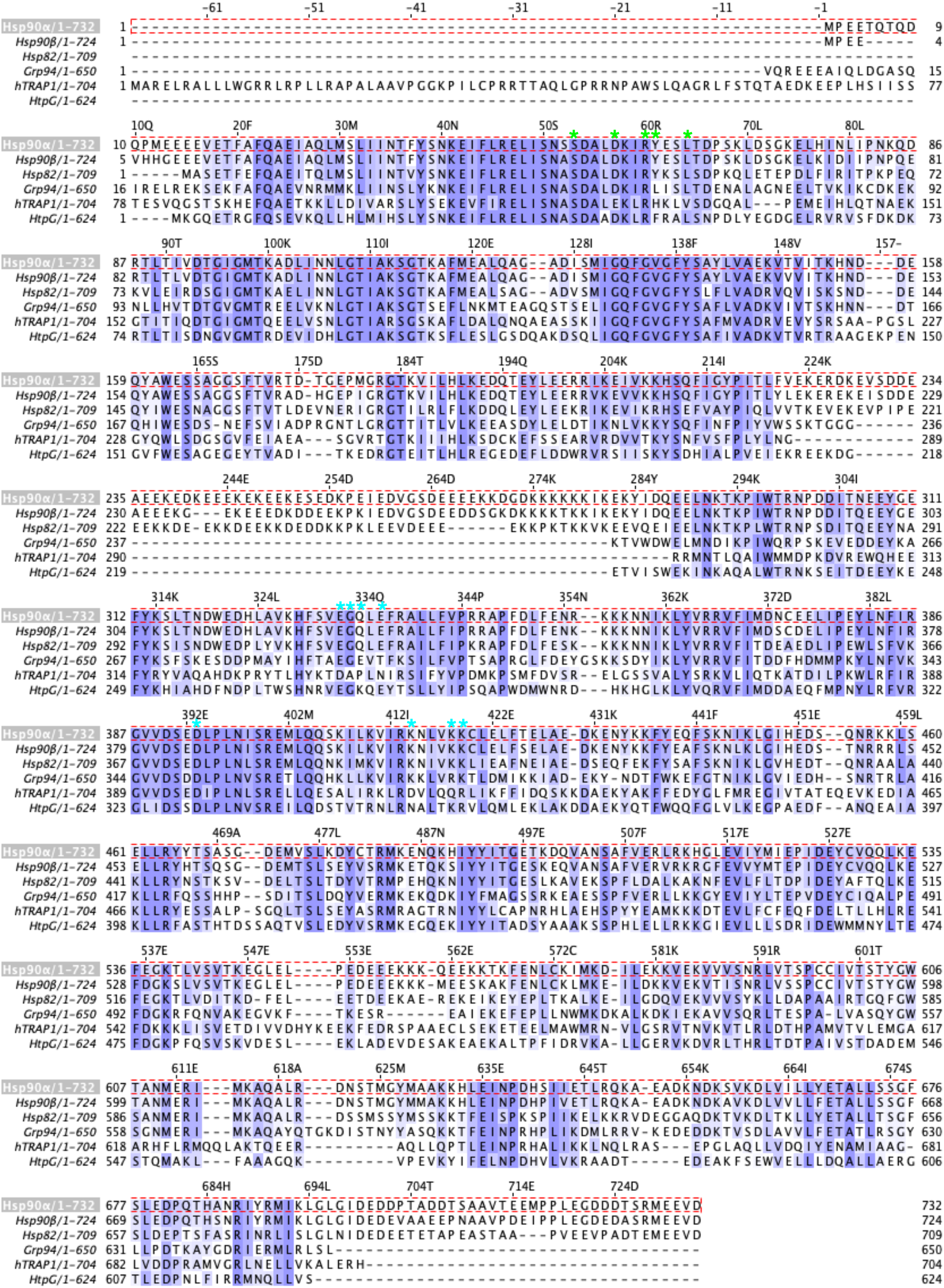
Sequence alignments of Hsp90 paralogs and homologs. Conserved residues are highlighted in purple. Key residues involved in Hsp90:Hsp70 interface I and II are marked with cyan and green asterisks, respectively. Hsp90α, human stress-induced; Hsp90β, human constitutively expressed; Hsp82, *S. cerevisiae* Hsp90; Grp94, human endoplasmic reticulum Hsp90; hTRAP1, mitochondrial Hsp90; HtpG*, E. coli* Hsp90. Multiple sequence alignments were generated using Clustal Omega and displayed using Jalview with BLOSUM62 Score color scheme.

**Extended Data Fig. 11.**
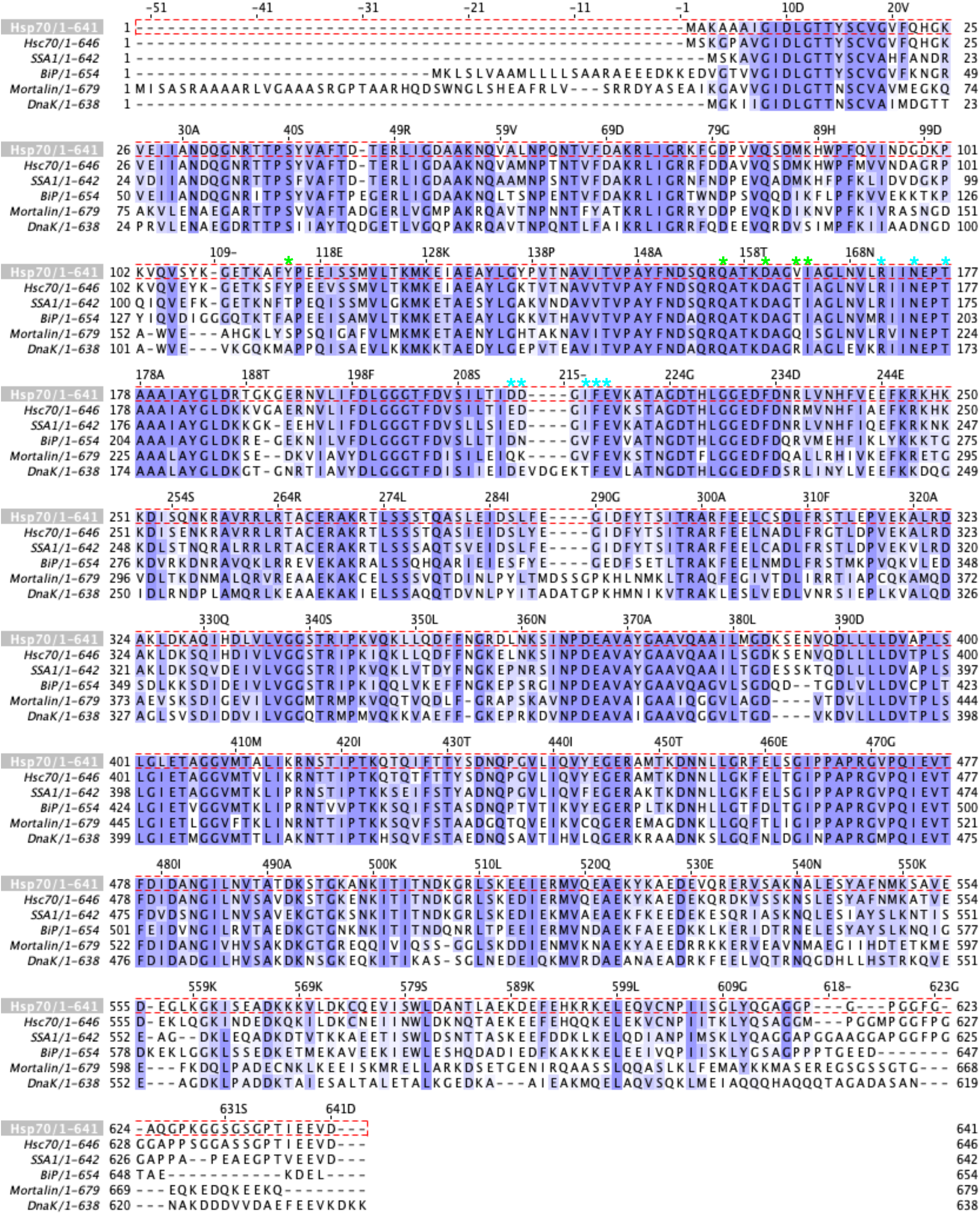
Sequence alignments of Hsp70 paralogs and homologs. Conserved residues are highlighted in purple. Key residues involved in Hsp90:Hsp70 interface I and II are marked with blue and red asterisks, respectively. Hsp70, human stress induced; Hsc70, human constitutively expressed; SSA1, *S. cerevisiae* Hsp70; BiP, human endoplasmic reticulum Hsp70; Mortalin, mitochondrial Hsp70; DnaK*, E. coli* Hsp70. Multiple sequence alignments were generated using Clustal Omega and displayed using Jalview with BLOSUM62 Score color scheme.

**Extended Data Fig. 12.**
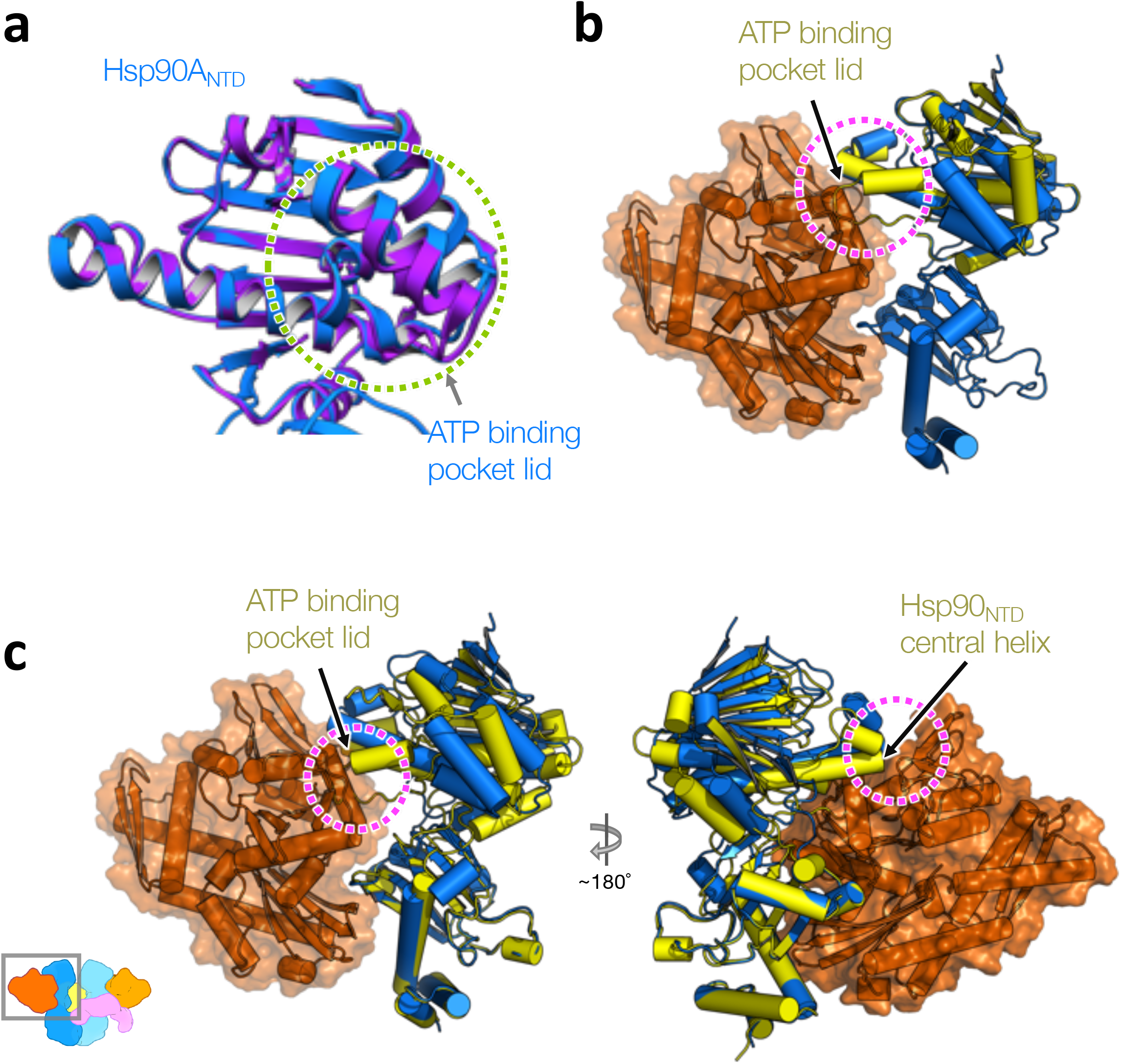
The Hsp90^ATP^ conformation is incompatible with the loading complex. **a**, Overlay of the crystal structure of Apo Hsp90 fragment (purple; PDB ID: 3t0h) to the Hsp90A_NTD_ (dark blue). Green circle highlights the open lid. **b**, Closure of the ATP pocket lid in the ATP state of Hsp90_NTD_ (the Hsp90α structure from the GR-maturation complex is in yellow, ribbon representation) clashes (magenta circle) with the Hsp70_NBD_ (orange, surface and ribbon representation) in the loading complex. The NTD fragment of Hsp90^ATP^ is aligned with the NTD of the Hsp90 in the loading complex. **c**, Superimposition of the ATP state of Hsp90^NTD-MD^ fragment (yellow) to the Hsp90A (dark blue) at the MD. Magenta circles indicate steric clashes of the ATP state of the Hsp90^NTD^ to the Hsp70^NBD^ (orange).

**Extended Data Fig. 13.**
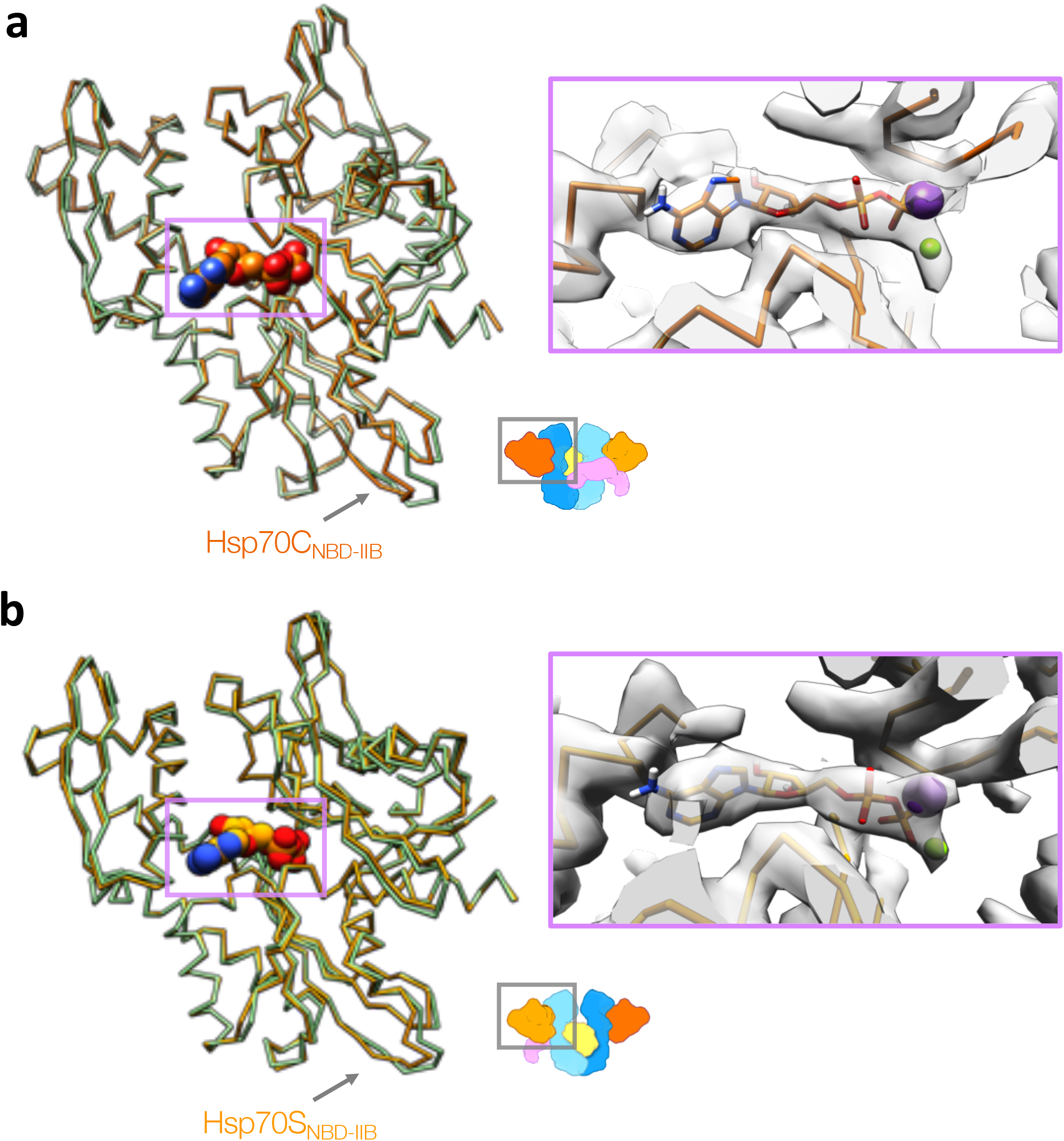
Marked deviation from the Hsp70_NBD-IIB_ to the crystal structure. **a,b,** Superposition of human ADP-bound crystal structure of Hsp70 (green; PDB ID: 3ay9) to the Hsp70C (**a**, dark orange) and Hsp70S (**b**, light orange) with backbone chain-trace representation. The purple rectangles highlight the bound ADPs (sphere representation) and the corresponding densities (right). Two metal ions were found in the Hsp70_NBD_. Based on the buffer compositions and the 3AT9 crystal structure, the two metal ion densities were assigned to be Potassium (purple sphere) and Magnesium (green sphere).

**Extended Data Fig. 14.**
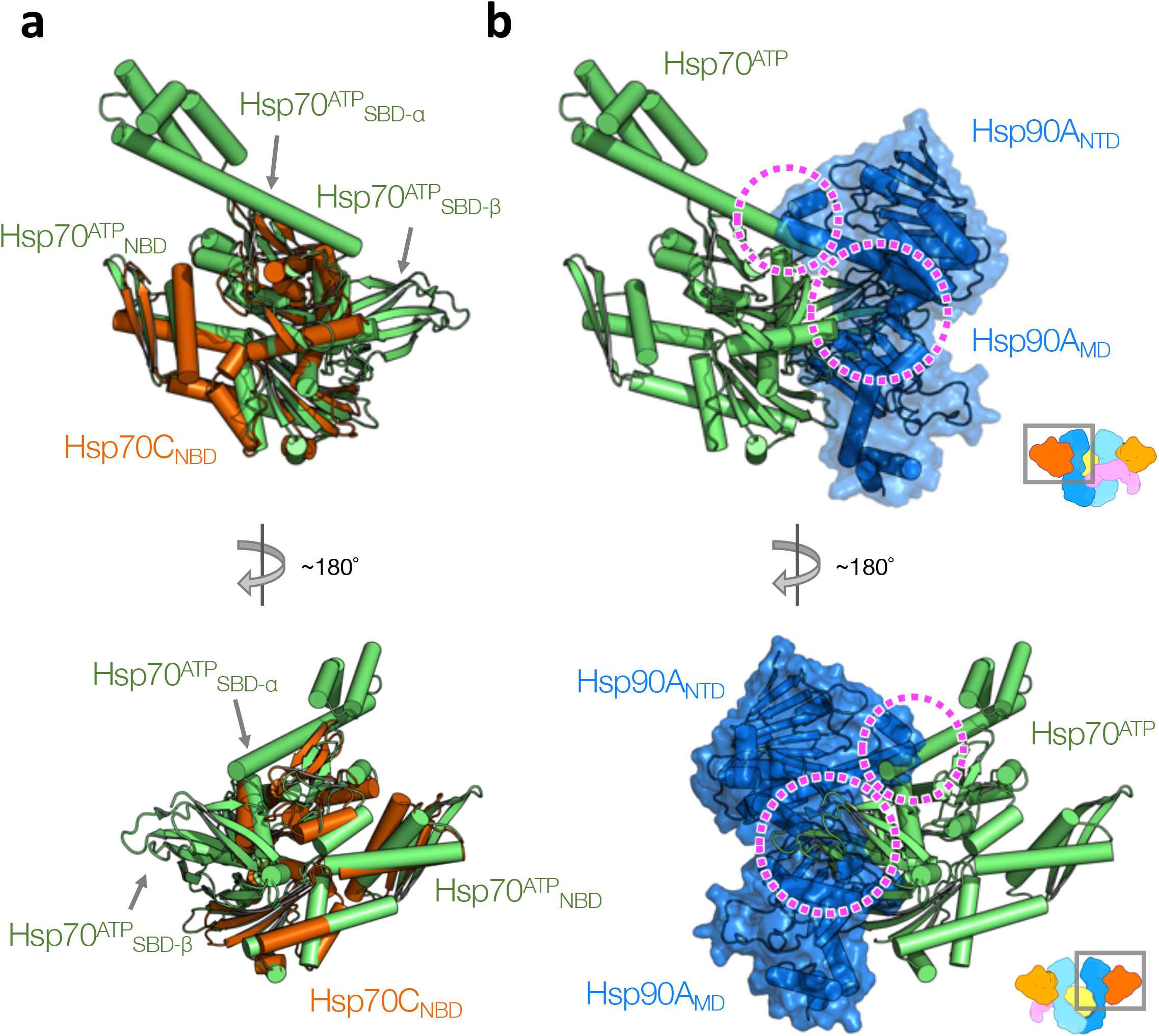
The Hsp70 ^ATP^ conformation is incompatible with the loading complex. **a**, Superposition of the Hsp70^ATP^ conformation (green; PDB ID: 4B9Q) to the Hsp70C_NBD_ (dark orange). Arrows indicate the two subdomains of Hsp70^ATP^ which cause serious steric clashes with the Hsp90 in the loading complex shown in (**b**). **b**, The superimposed Hsp70^ATP^ shown in (**a**) is fixed and the Hsp90 (dark blue; surface/ribbon representation) of the loading complex is present. Magenta circles highlight steric clashes caused by the two subdomains of Hsp70^ATP^_SBD_ (green) to the Hsp90A^NID-MD^.

**Extended Data Fig. 15.**
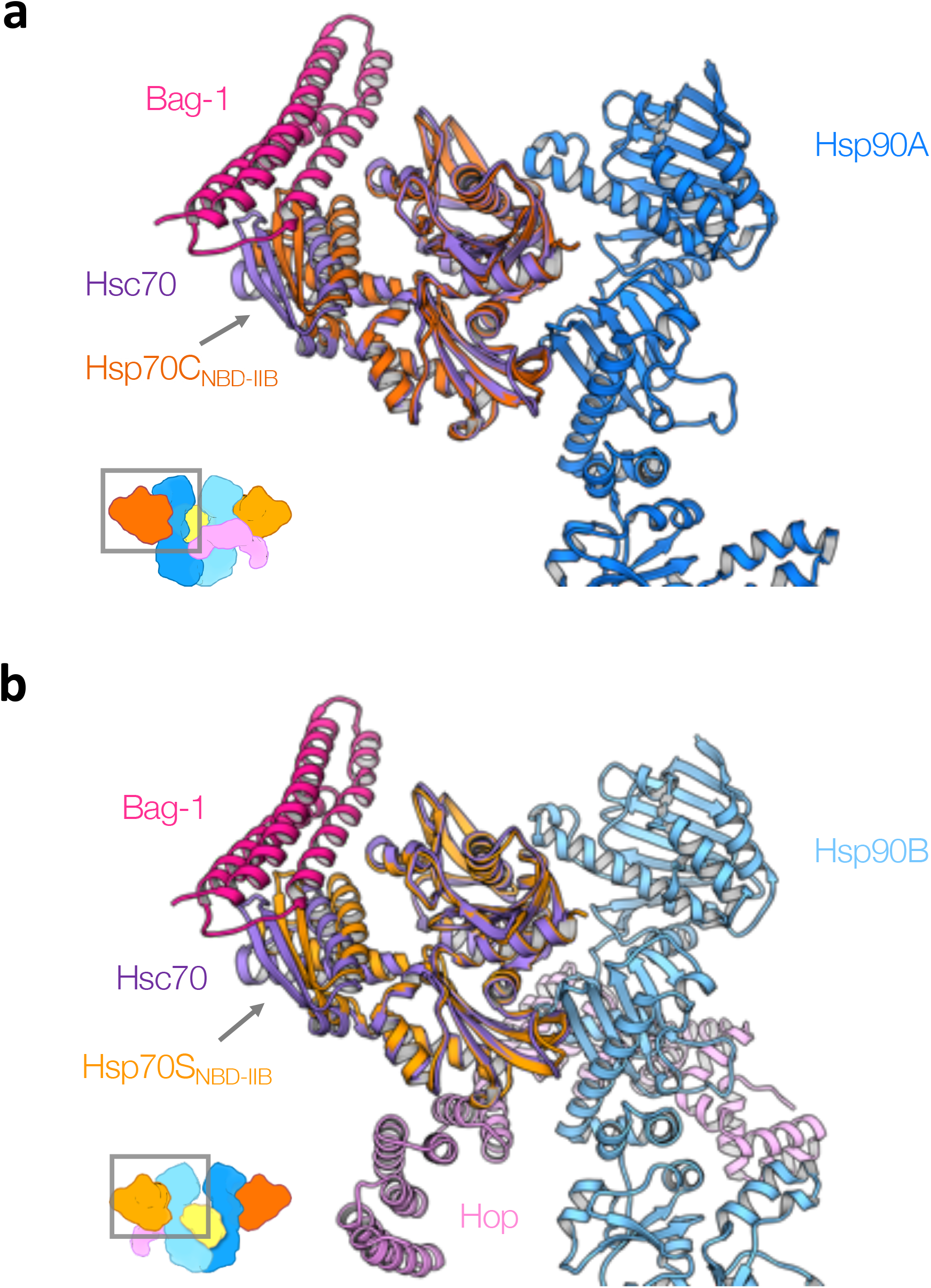
The canonical NEF binding sites are available in both of the Hsp70s. **a,b,** Superposition of the crystal structure (PDB ID: 1HX1) of the Bag-1:Hsc70 complex with the Hsp70C (**a**) and Hsp70S (**b**) on the loading complex. The Bag-1 and Hsc70 from the crystal structure are colored with magenta and purple, respectively. Components of the GR-loading complex are colored as in other figures and as labelled.

**Extended Data Fig. 16.**
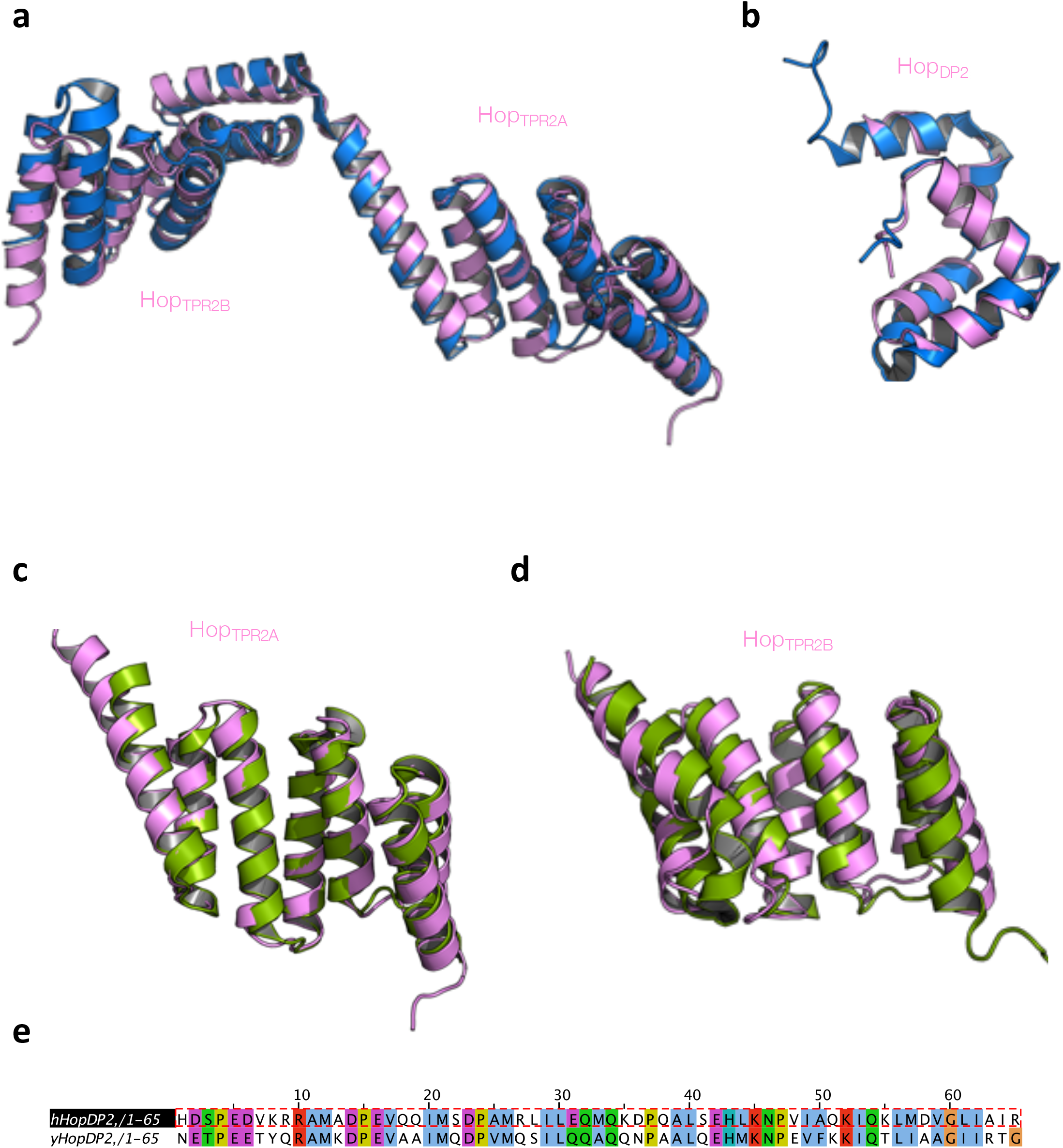
Comparison of the Hop_TPR2A-TPR2B-DP2_ structure with published experimental structures. **a**, Superposition of the Hop_TPR2A-TPR2B_ structure (pink) with the yeast crystal structure (blue; PDB ID: 3uq3). **b**, Superposition of the Hop_DP2_ (pink) with the yeast Hop_DP2_ NMR structure (blue; PDB ID: 2llw). **c**, Superposition of the Hop_TPR2A_ structure (pink) with the NMR structure (green; PDB ID: 2nc9). **d**, Superposition of the Hop_TPR2B_ structure (pink) with the NMR structure (green; PDB ID: 2lni) **e**, Sequence alignment used to derive the initial model for Hop_DP2_;the ClustalX color scheme available in Jalview was used to color the alignment.

**Extended Data Fig. 17.**
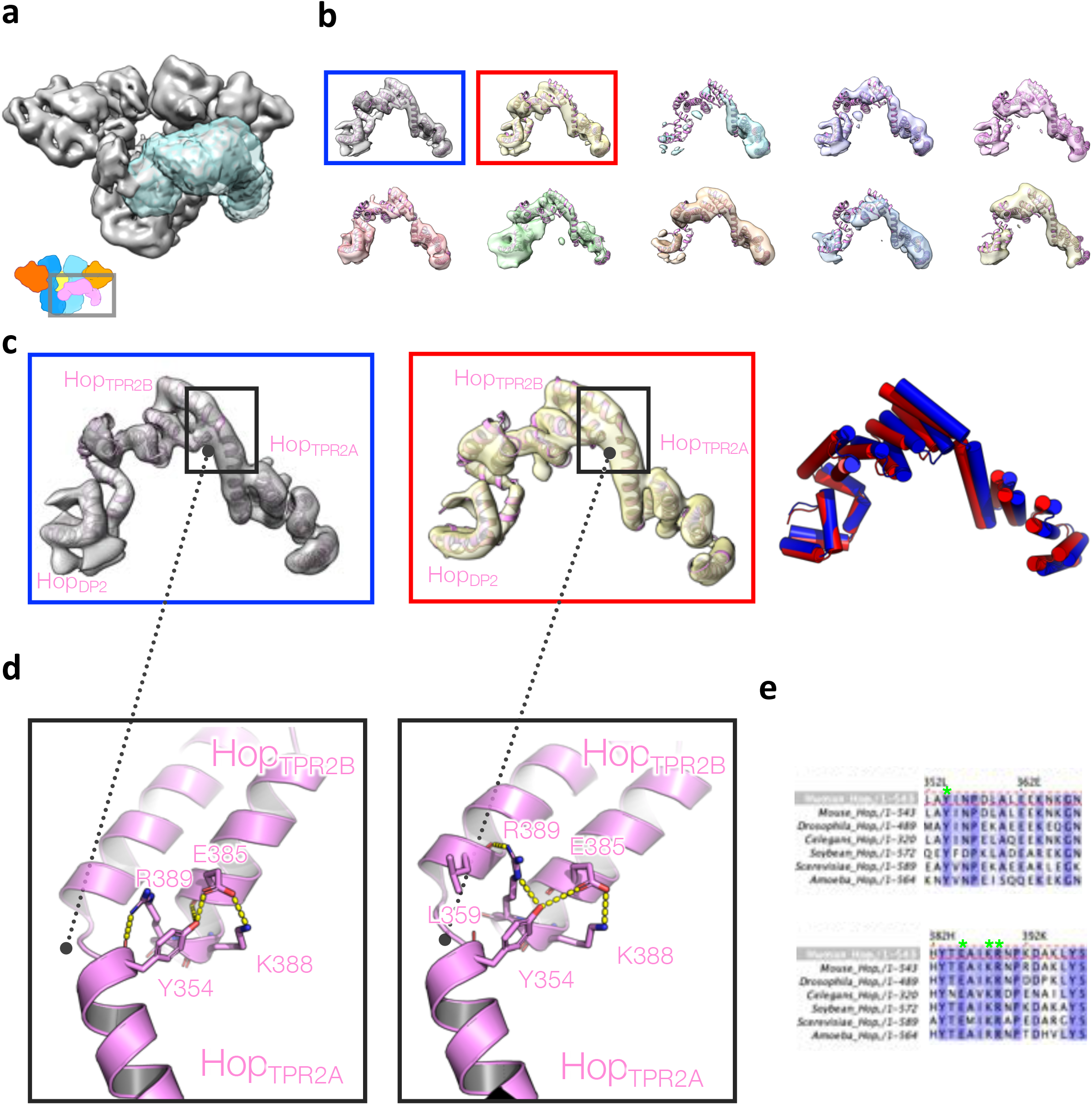
Conformation dynamics of Hop_TPR2A-TPR2B-DP2_ revealed by focused classification (without alignment). **a**, The mask used for the focused classification. **b**, 10 classes from RELION focused classification. Ribbon model (pink) fitted into the highest resolution class (blue rectangle) is kept static as a reference frame for the rest of the 9 maps. The top 2 high-resolution classes are highlighted with boxes. **c**, Models refined into the top 2 high-resolution classes (left and middle), respectively. Overlaid models without superposition (right) from the left and middle panels. **d**, A network of polar interaction defines the unique angle of the Hop_TPR2A_ and Hop_TPR2B_. This network is preserved in the two conformations. A similar network can also be found from the yeast homolog structure (PDB ID: 3uq3). **e**, Multiple sequence alignments of Hop from model systems. The residues involved in the network from (d) are indicated with green asterisks. Color scheme is BLOSUM62.

**Extended Data Fig. 18.**
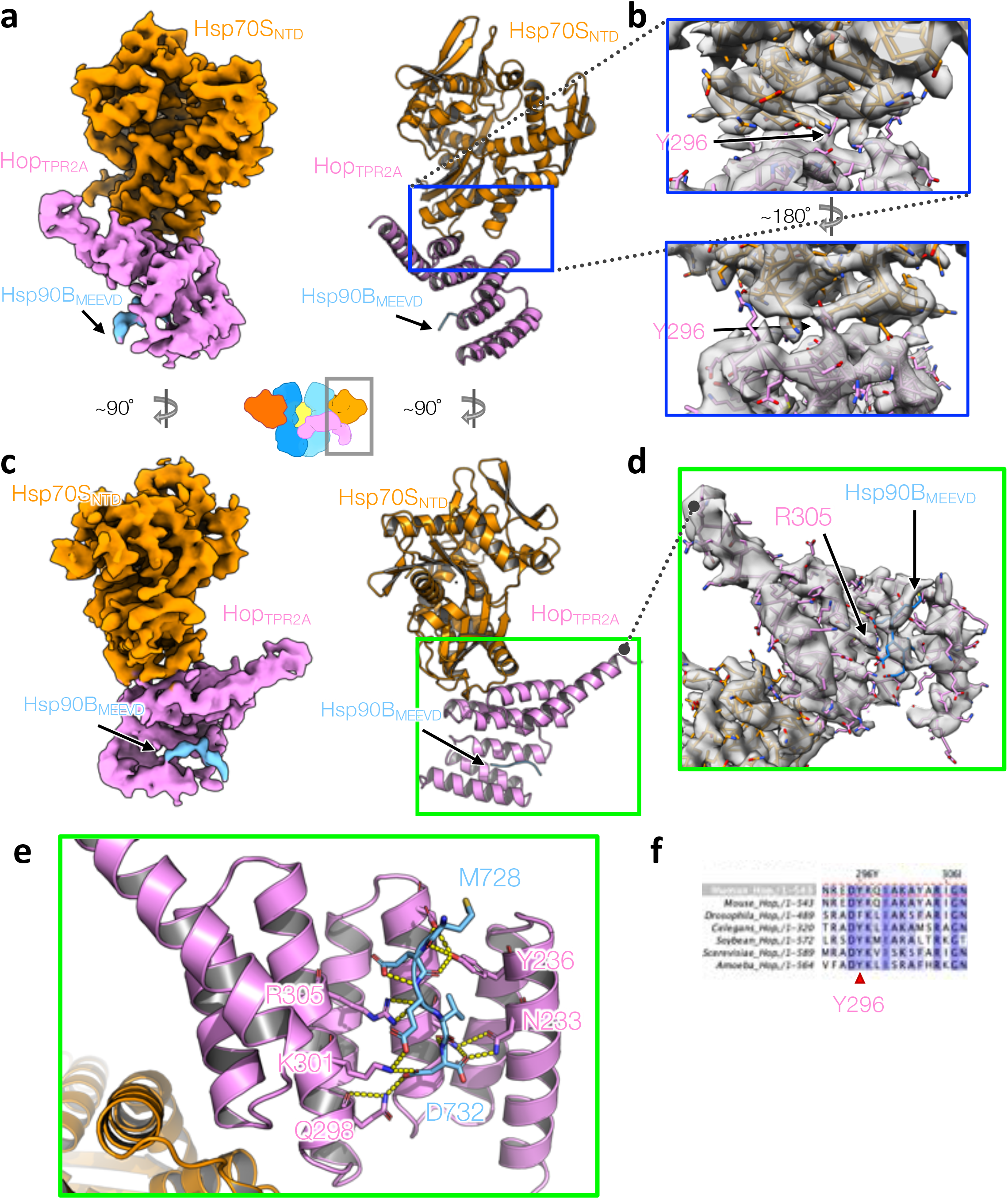
The atomic interactions of Hsp70S:Hop_TRP2A_:Hsp90B_MEEVD_ in the GR-loading complex. **a,c**, Left, the cryo-EM map from focused classification and refinement. Right, the atomic model with the corresponding views from the left. **b**, Close-up views of the Hsp70S_NTD_:Hop_TPR2A_ interface with the atomic model fit into the density. Sidechain density for Y296 from Hop_TPR2A_ is indicated with the arrow. **d**, Close-up view of the HopTPR2A. **e**, Close-up view of the atomic interactions of the MEEVD fragment from Hsp90B (light blue) and Hop_TPR2A_ (pink). Polar interactions are depicted with dashed lines. **f**, Sequence alignments of Hop indicate Y296 (red triangle) is highly conserved. Color scheme is BLOSUM62.

**Extended Data Fig. 19.**
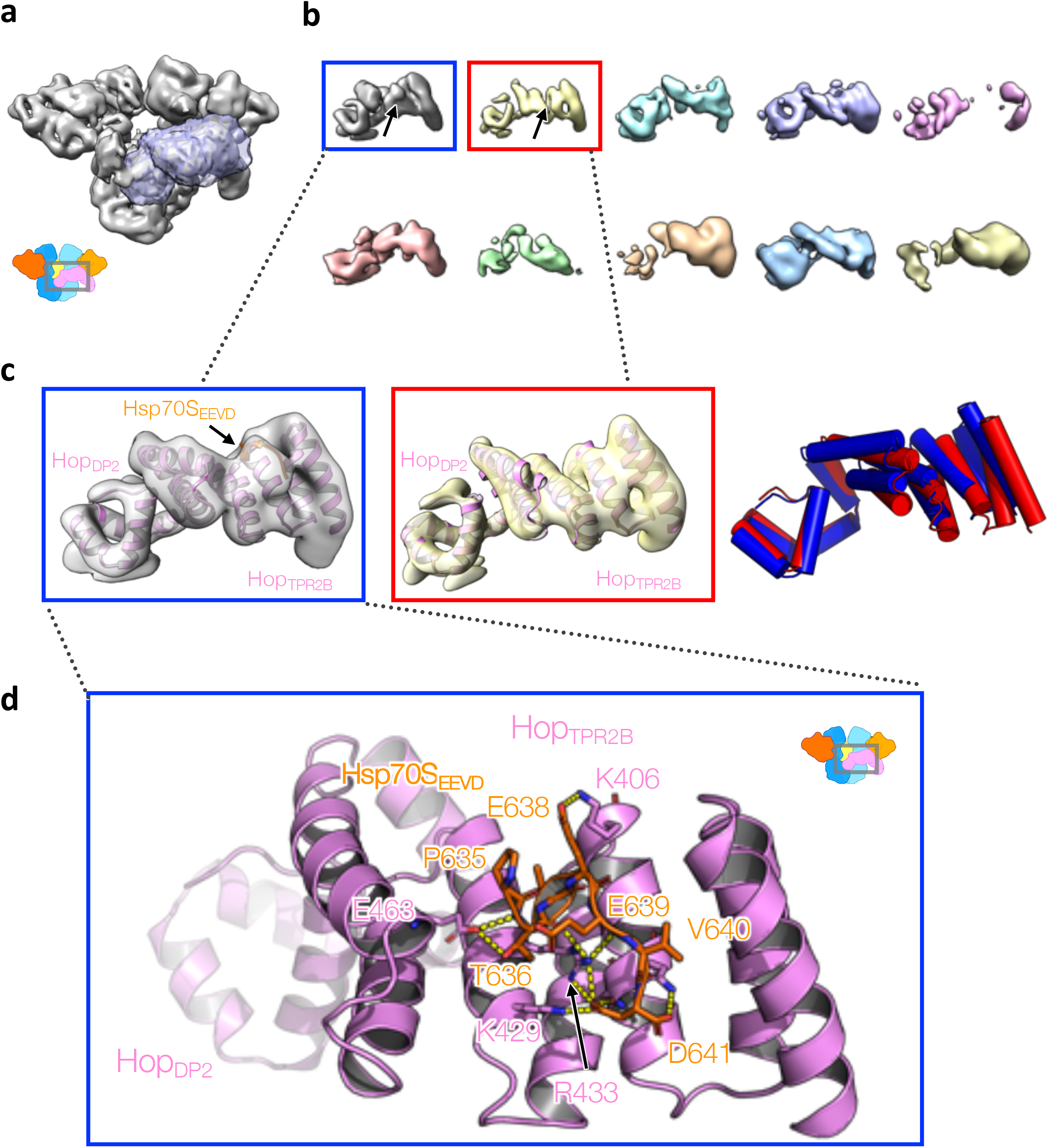
Hsp70S_EEVD_-bound Hop_TPR2B-DP2_ revealed by focused classification (without alignment). **a**, The mask used for the focused classification. **b**, The 10 classes from the focused classification. The top 2 high-resolution classes are highlighted with boxes. **c**, Models refined into the top 2 high-resolution classes (left and middle), respectively. Left, the highest resolution class has extra EEVD density (arrow) from Hsp70S, whereas the second-highest resolution class has no density in the EEVD binding pocket (middle panel). Right, overlaid models without superposition from the left and middle panels. **d**, Refined atomic model of Hop_TPR2B_ with Hsp70_EEVD_ fragment bound, in which the starting homology model was derived from the yeast homolog (PDB ID: 3UPV). Polar interactions are depicted with dashed lines.

**Extended Data Fig. 20.**
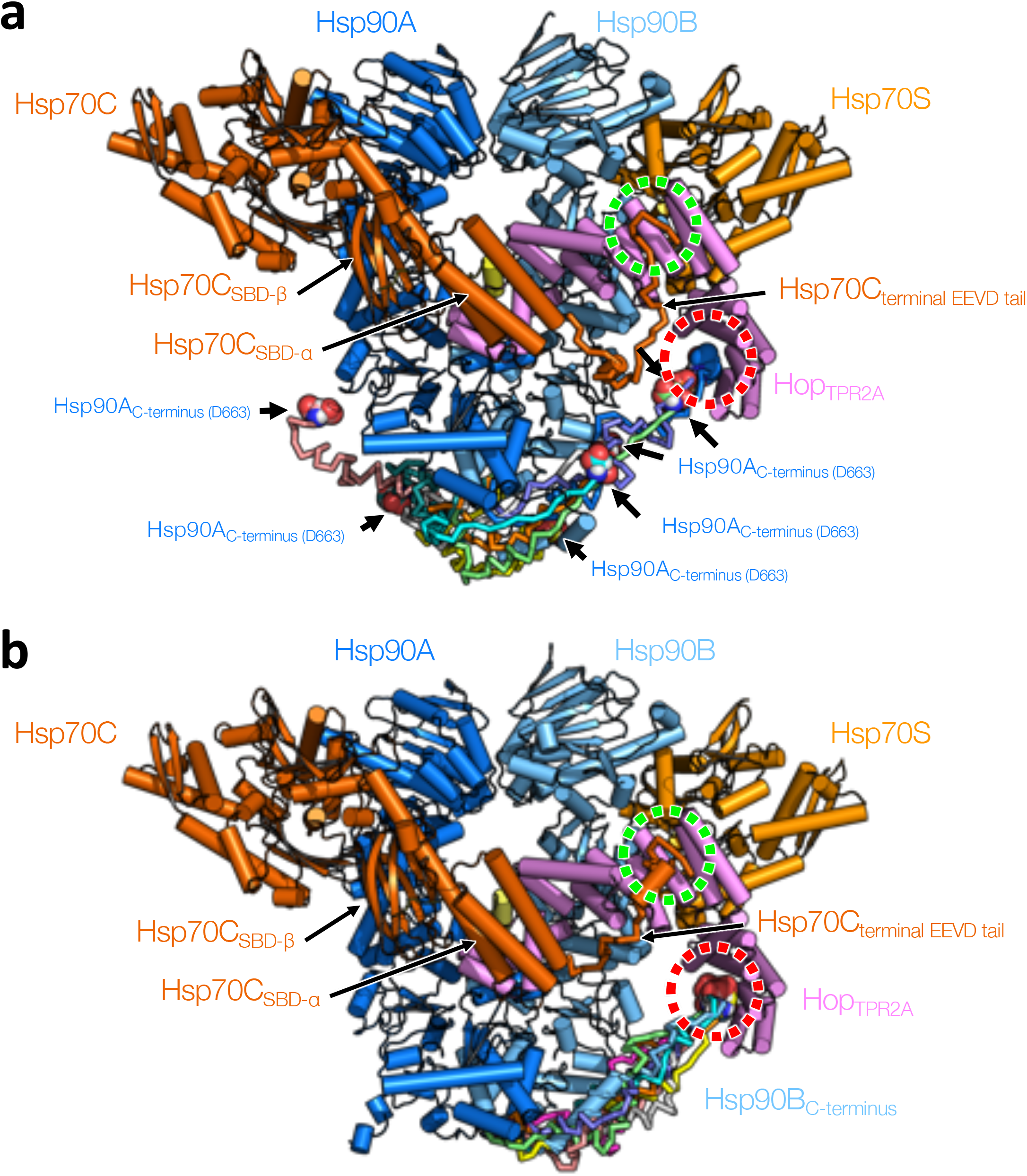
Modeling the EEVD tails from Hsp90/Hsp70 to Hop _TPR2A_ and Hop_TPR2B_, respectively. **a,b**, EEVD tail modeling starting with the loading complex model (shown in cartoon representation with the same color scheme) with a Hsp70/Hsp90 EEVD fragment bound to Hop _TPR2A_ (red dashed circle) and Hop_TPR2B_ (green dashed circle). Hsp70_SBD-α_ and its following ~30 tail was modeled to connect to the Hop_TPR2B_ EEVD-binding pocket. Two separate loop closure jobs were assigned to Hsp90A (top) and Hsp90B (bottom), respectively. The top 10 low-energy models are shown in ribbon representation with various colors and the very last residues of the MEEVD tail (D663) are shown in sphere representation with the black arrows pointed. Among the top 10 low-energy models, only 3 models from the Hsp90A job (top) successfully maintained the connection to Hop^TPR2A^ (red circle), whereas all 10 models from the Hsp90B job (bottom) were able to maintain the binding to Hop^TPR2A^ (red circle).

**Extended Data Fig. 21.**
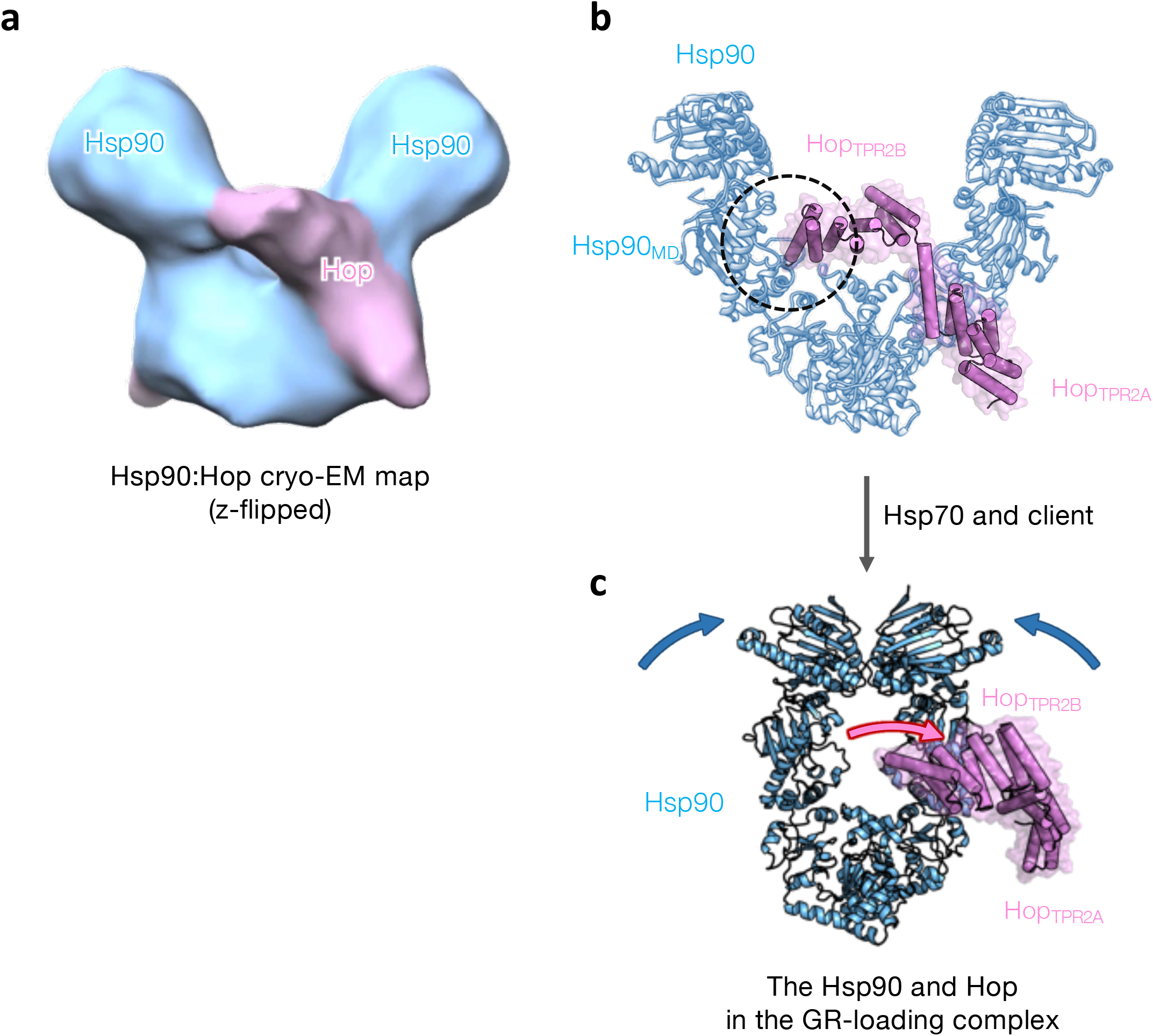
Hop may first prepare Hsp90 for Hsp70 and client interaction. **a**, A mirror image of the ~15-Å-resolution cryo-EM map of the Hsp90:Hop complex (Southworth & Agard, 2014). In cryo-EM single-particle analysis, there is a 50% chance to get a mirror reconstruction and the correct handedness is difficult to distinguish in medium to low resolution. **b**, Guided by the flipped map and the GR-loading complex structure, now the Hop density (pink shade in (**a**)) can be interpreted as the TPR2A-TPR2B module of Hop as it possesses a unique dumb-bell-like structure. Dashed circle indicates a contact of Hop_TPR2B_ to Hsp90_MD_ in this state. **c**, The Hsp90 and Hop structure in the GR-loading complex. After interacting with Hsp70 and GR, the Hsp90 dimer and Hop_TPR2A-TPR2B_ undergo conformational changes from the Hsp90:Hop complex, which are indicated with the blue and pink arrows, respectively.

**Extended Data Fig. 22.**
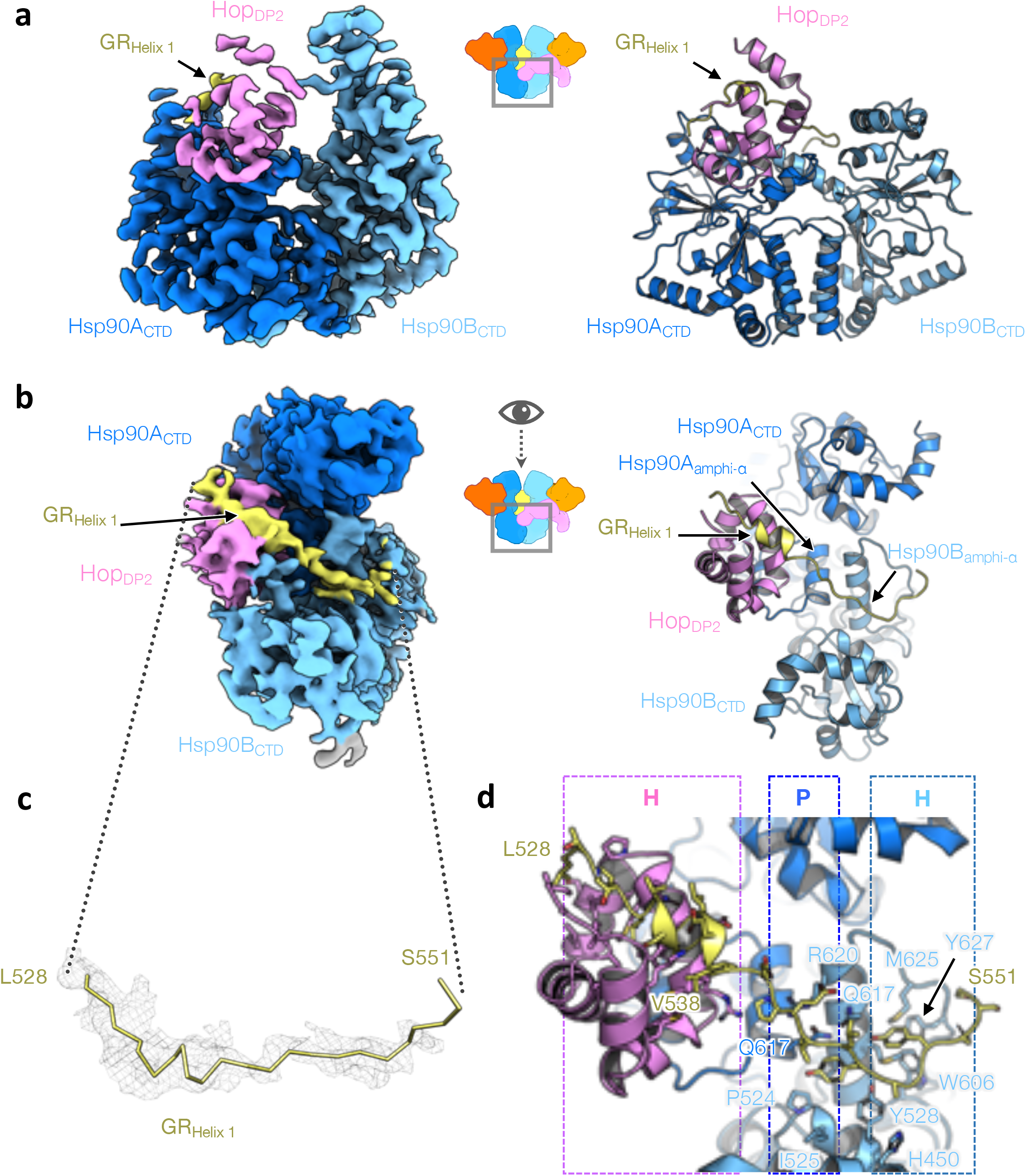
GR Helix 1 motif’s interactions with Hsp90 and Hop_DP2_. **a**, The focused map of the Hsp90AB_CTD_:Hop_DP2_:GR_N-term_(left) and the atomic model shown in representation (right). **b**, The top view of the reconstruction and model shown in (**a**). **c**, The density (mesh) for the GR_Helix 1_ motif (residues 528-551) gripped by Hsp90 and Hop_DP2_. **d**, The atomic interactions of the GR_Helix 1_ motif with Hsp90 and Hop_DP2_. Residues in contact with the GR motif are shown in stick representation. The types of molecular interaction Hsp90 and Hop_DP2_ provide are indicated on the top, where H and P denote hydrophobic and polar interactions, respectively.

**Extended Data Fig. 23.**
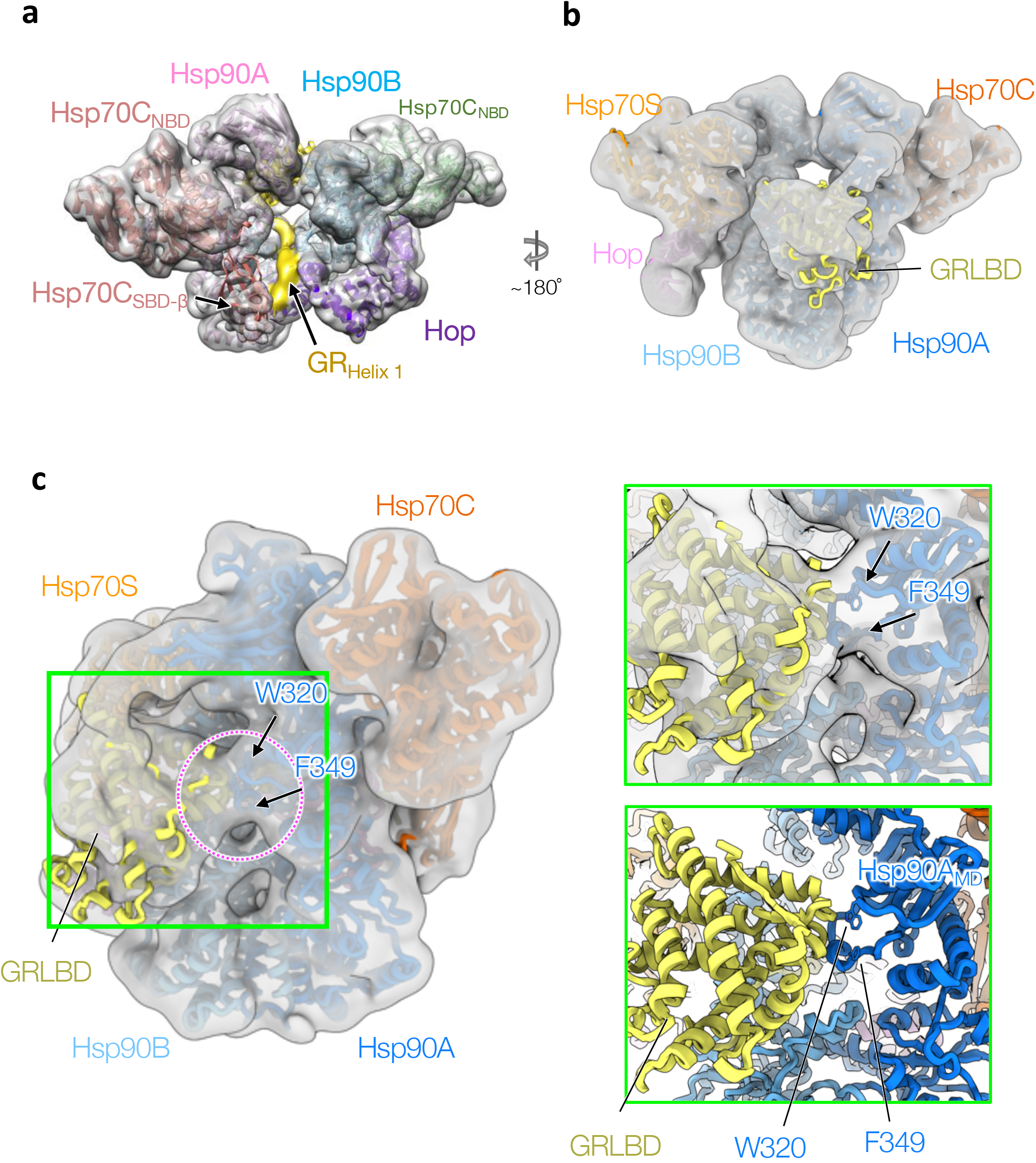
GR is unfolded, threaded through the Hsp90 lumen and bound by Hsp90, Hop and Hsp70 density. **a**, A lumen density of GR connects to the globular part of GR on the other side of Hsp90. **b**, Docking of the GRLBD to the low-pass filtered map shows that the low-resolution GR density can fit the rest of the GRLBD. **c**, The low-pass filtered map shows that W320 and F349 (arrows) of Hsp90A in the loading complex are in contact with GRLBD.

**Extended Data Fig. 24.**
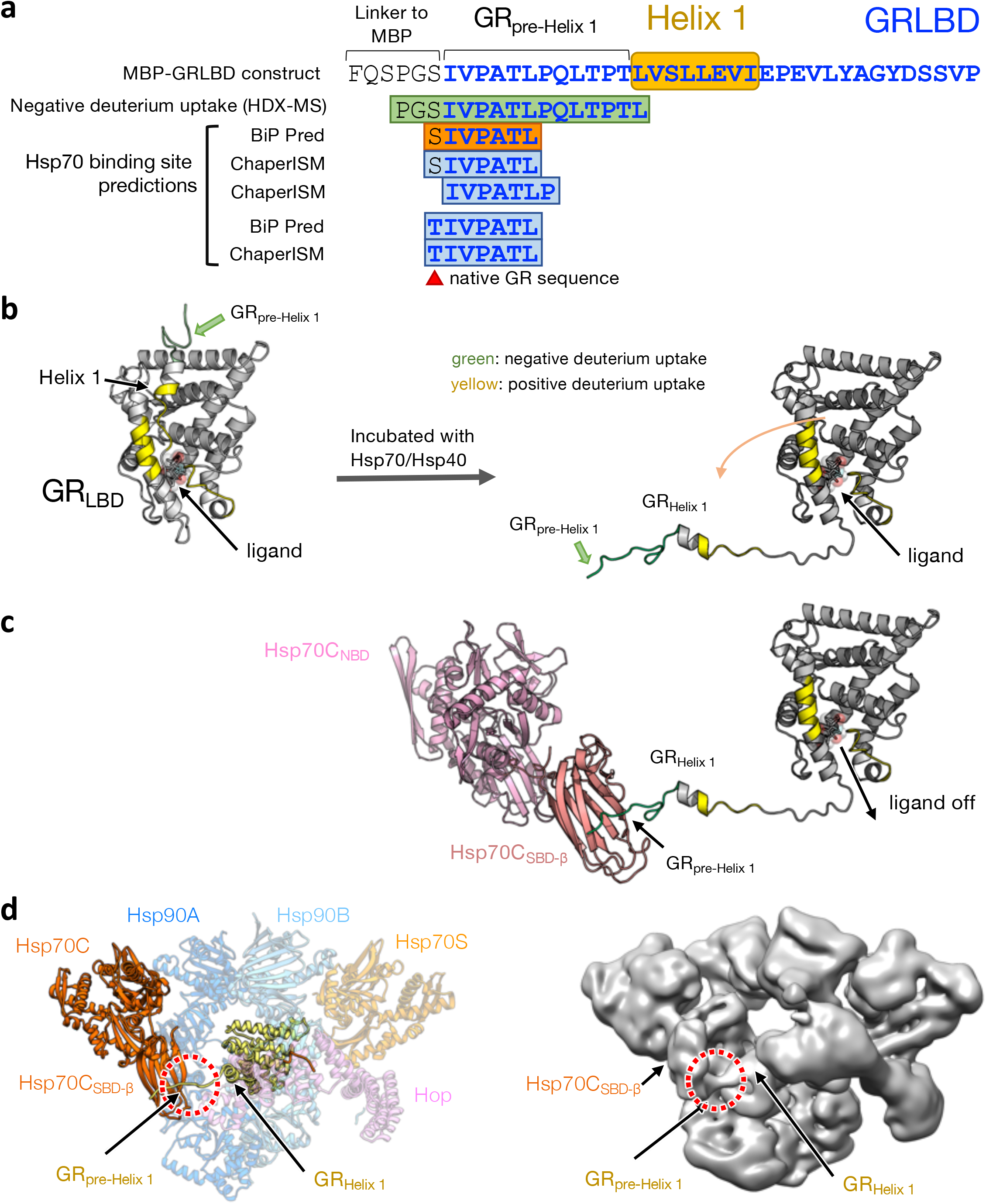
Hsp70 inhibits GR by binding the pre-Helix 1 region of GR. **a**, After engaging with Hsp70/Hsp40, the GR_pte-Helix 1_ region exhibits negative deuterium uptake in a HD-exchange mass spectrometry (HDX-MS) experiment (Krschke et al., 2014). In the GR_pre-Helix 1_ region, there are Hsp70 binding sites predicted by two state-of-the-art algorithms (BiP Pred and ChaperISM). **b**, Left, GRLBD crystal structure (1 m2z) colored by the change of deuterium uptake (HDX-MS was retrieved from Kirschke et al., 2014); green: negative; yellow:positive. Right, the detachment of the entire GR Helix 1 motif explains the positive deuterium uptake around the ligand-binding pocket. **c**, The negative uptake of the pre-Helix region can be explained by binding of Hsp70. Together (**b**) and (**c**) provide a molecular mechanism describing how Hsp70 inhibits GR ligand binding. **d**, GR’s pre-Helix 1 remains bound to Hsp70 (red circles) in loading complex. Left, atomic model with ribbon presentation. Right, a low-pass filtered cryo-EM map of the loading complex.

**Extended Data Fig. 25.**
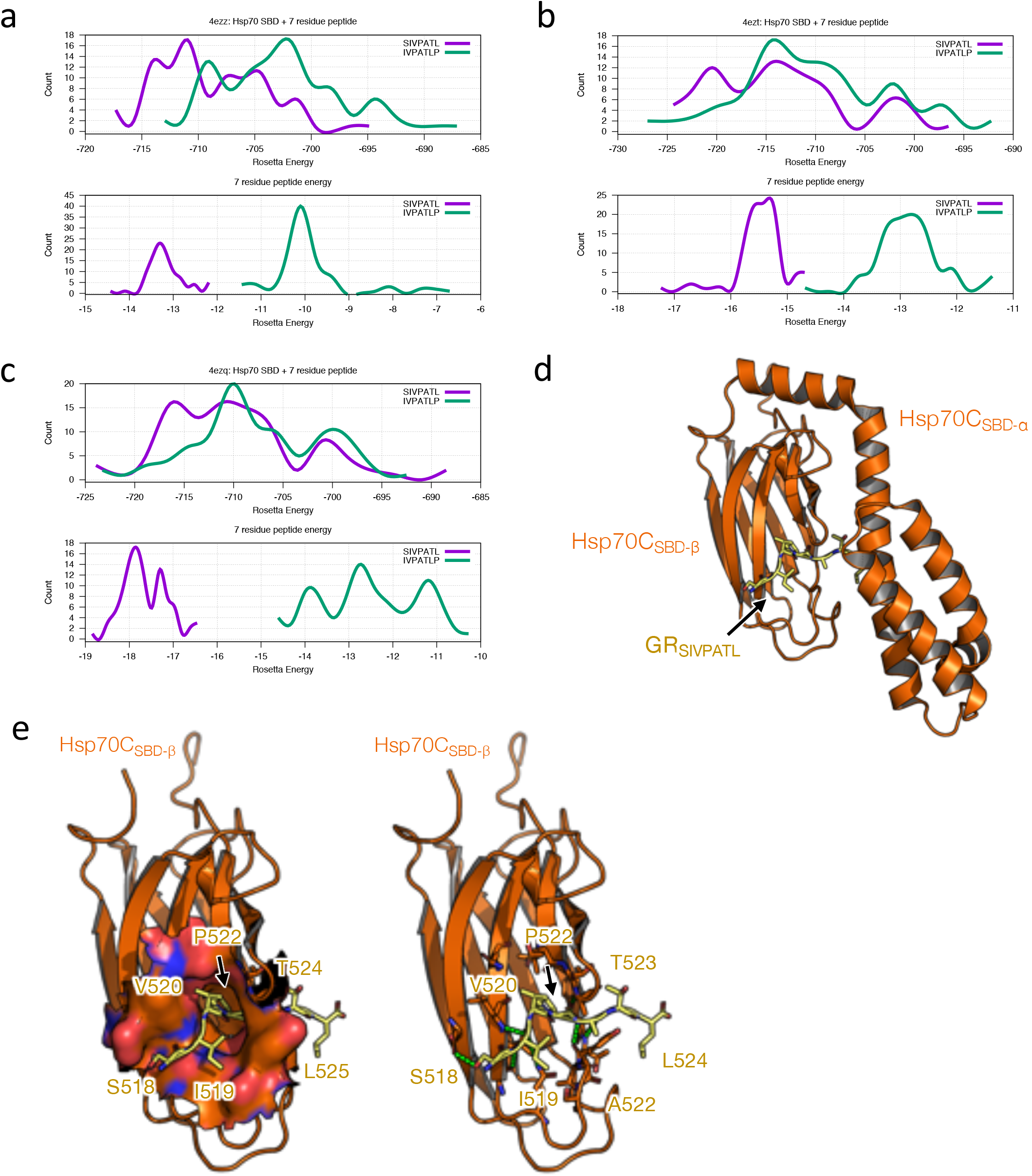
Determination of the GR segment that Hsp70C_SBD_ binds. **a,** Two 7-residue GR segments (SIVPATL and IVPATLP) of a continuous sequence (residues 518-525; note that residue 518 in the native GR sequence is T, not S) in the pre-Helix 1 regions are predicted as Hsp70 binding sites. Using the crystal structures of *E. coli* Hsp70_SBD_ (DnaK) with “reverse” binding peptides as templates (PDB ID: 4ezz (**a**), 4ezt (**b**), and 4ezq (**c**)), sequences of the two GR segment and human Hsp70 were threaded to the backbone of the crystal structures, resulting in two starting homology models of Hsp70_SBD_:peptide for each template. Rosetta full-atom energy minimization was carried out for the individual threaded models. 100 models were generated for each threaded model. Density plots show the energy distribution (Y-axis=count) of the 100 models, where the top plot shows the total energy (X-axis) from Hsp70:peptide and the bottom plot shows only energy contributed by the peptide. **b**, Same as (**a**), but using 4ezt as the template. **c**, Same as (**b**), but using 4ezq as the template. **d**, The selected model, which was the lowest energy model from the job using 4ezq as the template and SIVPATL as the bound peptide. **e**, Closeup view of the Hsp70_SBD_ from (**d**) with surface representation with positive/negative charged residues colored in blue/red (left) and stick representation (right) around the peptide binding pocket. Right, green dashed lines indicate polar interactions.

**Extended Data Fig. 26.**
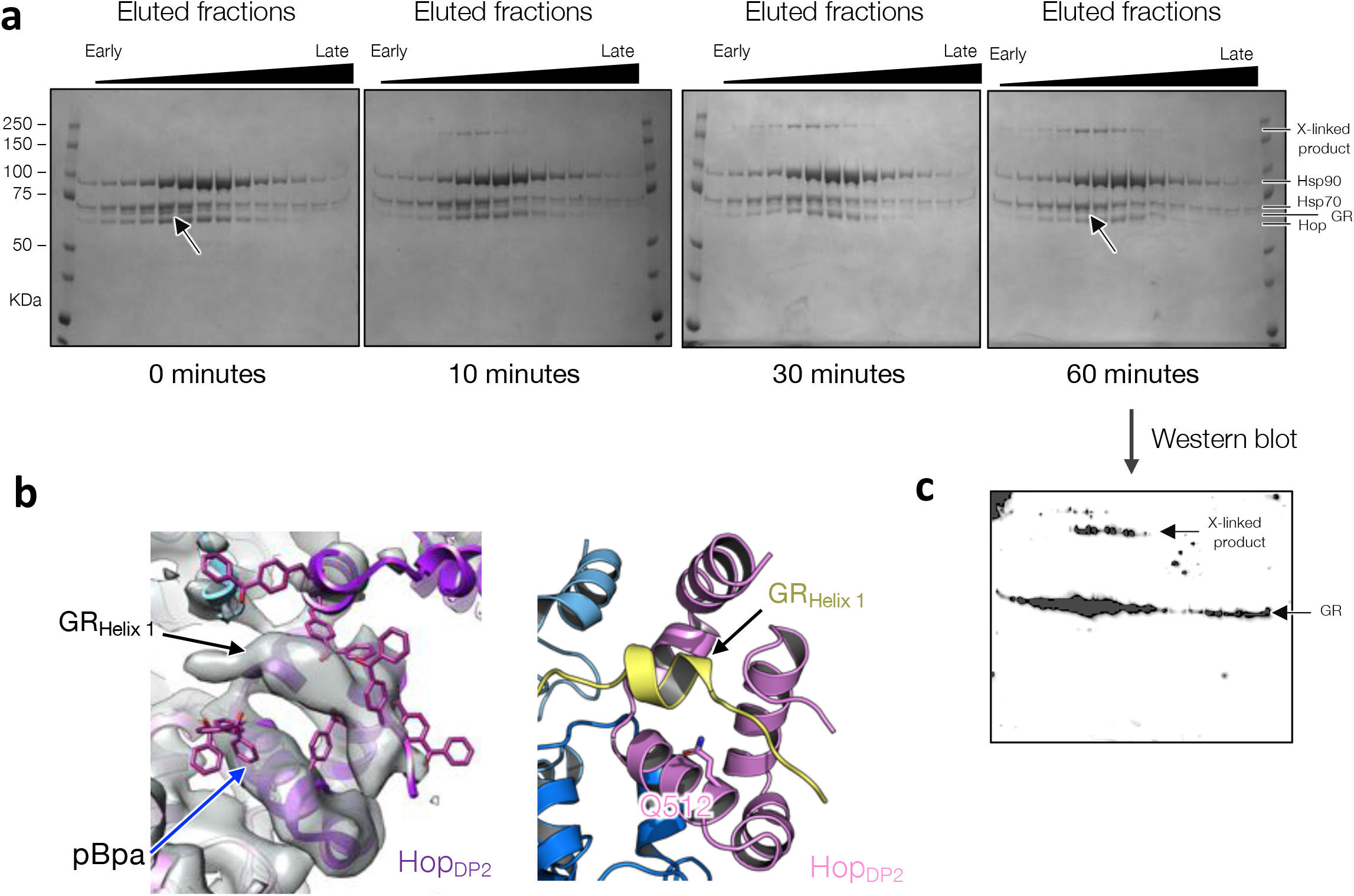
The density Hop_DP2_ binds belongs to GR. **a**, A time course of UV-exposed GR-loading complex analyzed by SDS-PAGE and visualized by Coomassie staining. Whole fractions of GR-loading complex eluted from the size-exclusion column were exposed to UV using a gel imager. Arrows at 0 and 60 minutes indicate a reduced intensity of the GR band over the time course. **b**, Modeling the photoreactive crosslinker p-benzoyl-L-phenylalanine (pBpa) (magenta) on various positions of Hop_DP2_. The blue arrow on the left panel points at the selected position, Q512 (right panel). **c**, Western blot of the SDS-PAGE gel after a 60-minute UV exposure, using anti-MBP antibody to detect the MBP-tagged GR.

**Extended Data Fig. 27.**
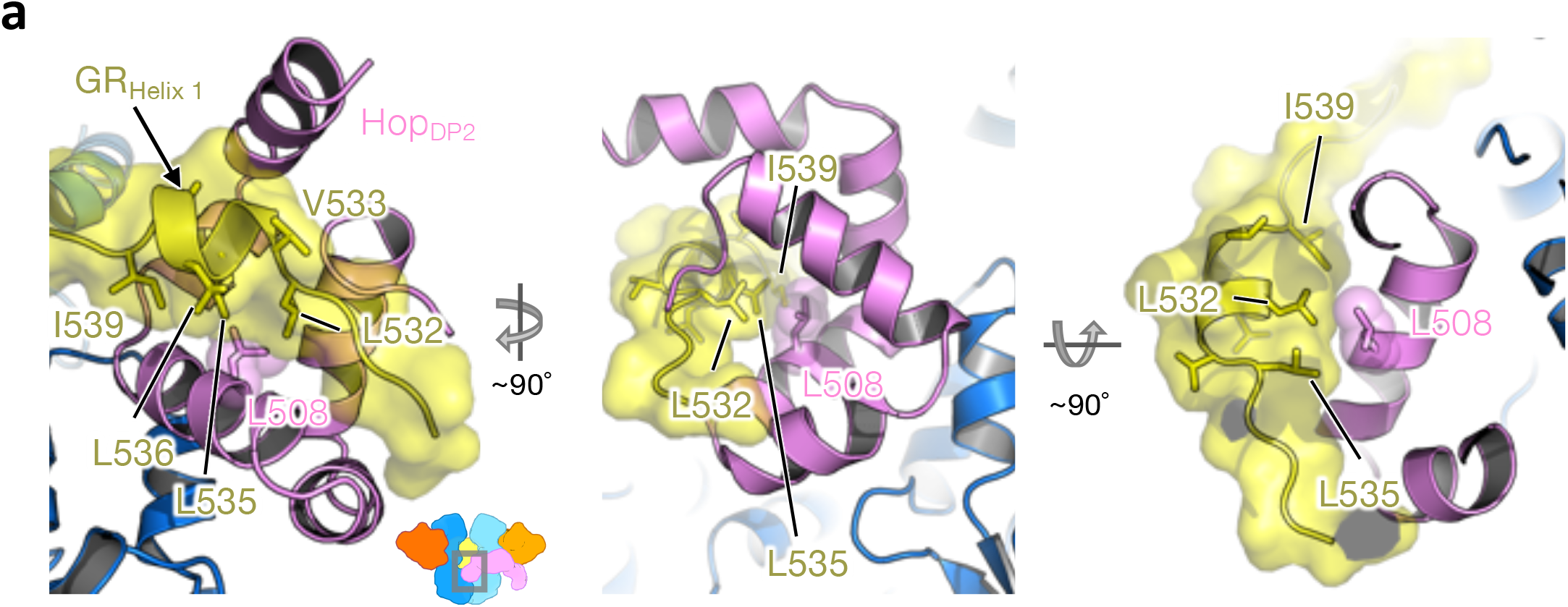
Molecular basis for the effect of the HOP L508A mutation, which completely abrogates GR function in vivo. **a**, L508 is located on the hydrophobic palm of DP2, interacting closely with the LXXLL motif of GR_Helix1_ through hydrophobic interactions (left, middle and right).

**Extended Data Fig. 28.**
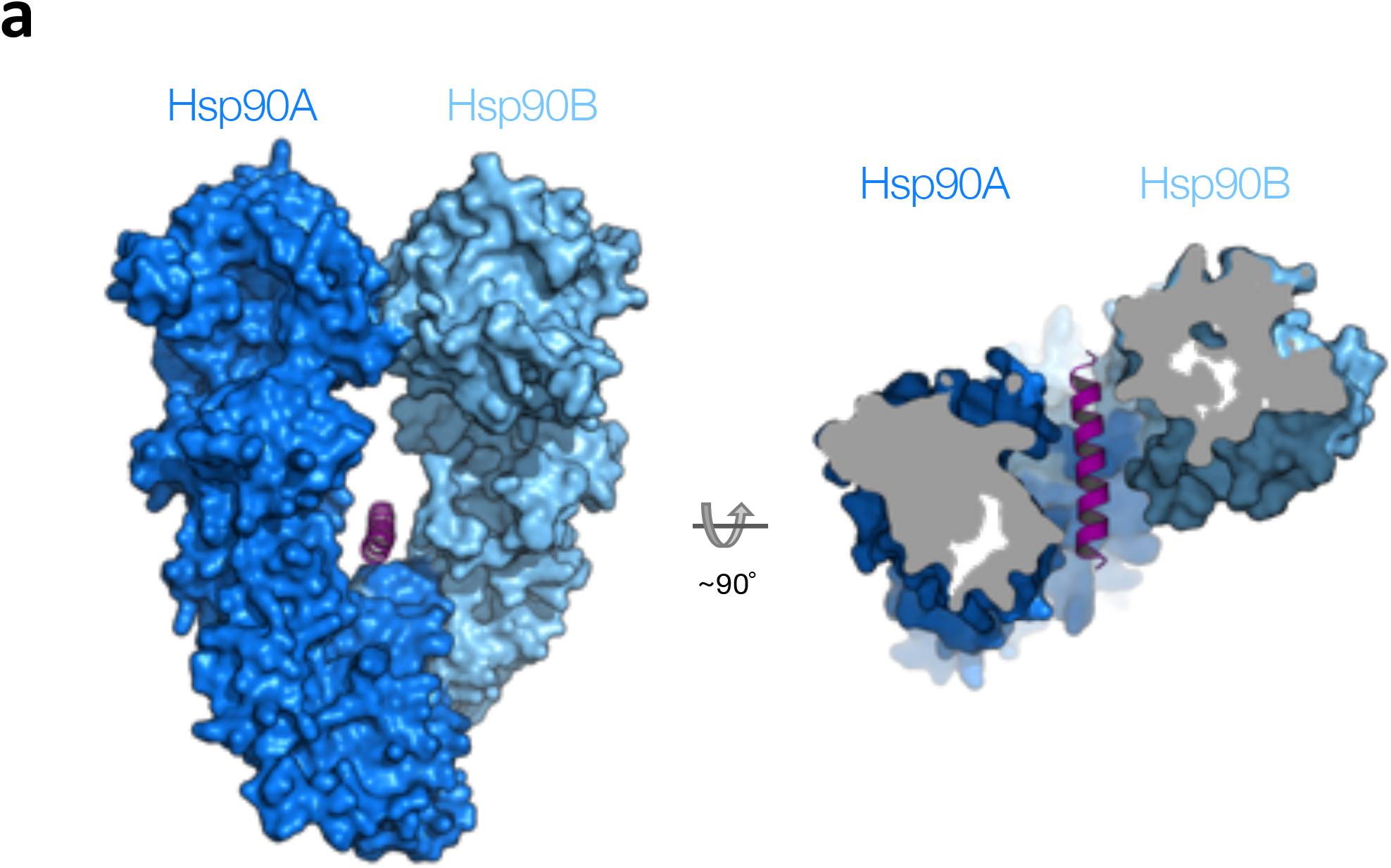
The lumen of the semi-closed Hsp90 presented in the loading complex can fit a helix. **a**, A helix (magenta) can be accommodated in the semi-closed Hsp90. Front view (left) and top view (right).

**Extended Data Table 1.** Cryo-EM data collection

| Data set | D93N-I | D93N-II | D93N-III | D93N-IV |
| --- | --- | --- | --- | --- |
| Microscope | Titan Krios | Titan Krios | Titan Krios | Titan Krios |
| Camera | K2 Summit | K2 Summit | K2 Summit | K2 Summit |
| **Data collection and processing** |  |  |  |  |
| Magnification | 22,500 | 22,500 | 22,500 | 22,500 |
| Voltage (kV) | 300 | 300 | 300 | 300 |
| Electron Exposure (e<sup>-</sup>/Å<sup>2</sup>) | 80 | 80 | 70 | 70 |
| Exposure time (sec) | 12 | 16 | 14 | 14 |
| Defocus range (μm) | -0.5 to -2.3 | -0.5 to -2.5 | -1.0 to -2.5 | -0.5 to -2.5 |
| Pixel Size (Å/pixel) | 1.059 | 1.059 | 1.059 | 1.059 |
| Micrographs collected (no.) | 3145 | 3213 | 489 | 4248 |
| Particles picks (no.) | 986,519 | 1,397,986* |  | 1,728,005 |
| Refined particles (no.) | 139,412 | 214,315* |  | 282,329 |

\* Data sets II and III were merged and processed together.

